# ATRX guards against aberrant differentiation in mesenchymal progenitor cells

**DOI:** 10.1101/2023.08.08.552433

**Authors:** Yan Fang, Douglas Barrows, Yakshi Dabas, Thomas S Carroll, William D. Tap, Benjamin A. Nacev

**Affiliations:** Department of Medicine, Memorial Sloan Kettering Cancer Center, New York, NY10065; Laboratory of Chromatin Biology and Epigenetics, The Rockefeller University, New York, NY10065; Bioinformatics Resource Center, The Rockefeller University, New York, NY10065; Department of Medicine, University of Pittsburgh, Pittsburgh, PA 15213; Department of Pathology, University of Pittsburgh, Pittsburgh, PA 15213; UPMC Hillman Cancer Center, Pittsburgh, PA 15213

## Abstract

Alterations in the tumor suppressor *ATRX* are recurrently observed in several cancer types including sarcomas, which are mesenchymal neoplasms. ATRX has multiple epigenetic functions including heterochromatin formation and maintenance and regulation of transcription through modulation of chromatin accessibility. Here, we show in murine mesenchymal progenitor cells (MPCs) that *Atrx* deficiency aberrantly activated mesenchymal differentiation programs. This includes adipogenic pathways where ATRX loss induced expression of adipogenic transcription factors (*Pparγ* and *Cebpα*) and enhanced adipogenic differentiation in response to differentiation stimuli. These changes are linked to loss of heterochromatin near mesenchymal lineage genes together with increased chromatin accessibility and gains of active chromatin marks at putative enhancer elements and promoters. Finally, we observed depletion of H3K9me3 at transposable elements, which are derepressed including near mesenchymal genes where they could serve as regulatory elements. Our results demonstrate that ATRX functions to buffer against differentiation in mesenchymal progenitor cells, which has implications for understanding ATRX loss of function in sarcomas.

## Introduction

Alpha Thalassemia/Mental Retardation Syndrome X-Linked (ATRX) belongs to the SWI/SNF family of ATP-dependent chromatin remodeling proteins [1]. Germline mutations in ATRX cause cognitive impairment as part of the alpha-thalassemia (ATR-X) syndrome, which is accompanied by disturbances of DNA methylation [2], highlighting the function of ATRX in development [3]. Somatic alterations in *ATRX* occur in cancers, such as sarcomas [4].

ATRX is an important epigenetic regulator, functioning as a chromatin remodeler and promoting the formation and maintenance of heterochromatin [5]. For example, ATRX cooperates with the H3K9 methyltransferase SETDB1 to establish and maintain heterochromatin, including at retrotransposons [5]. ATRX also binds to H3K9me3 via its ADD domain to directly target it to heterochromatin, which may be important for its role in maintaining H3K9me3 domains [5]. ATRX also binds to DAXX, functioning as a histone chaperone depositing H3.3 into telomeric regions [6].

Loss of function genetic alterations in *ATRX* are highly recurrent in several cancers including gliomas, pancreatic neuroendocrine tumors, and multiple sarcoma subtypes [7, 8]. Within sarcomas, *ATRX* is altered in more than 10 percent of undifferentiated pleomorphic sarcoma, leiomyosarcoma, myxofibrosarcoma, perivascular epithelioid tumors, pleomorphic liposarcoma and angiosarcoma. In uterine leiomyosarcoma (ULMS), the frequency of *ATRX* alterations is approximately one third of cases [7]. Consistent with these genetic studies, loss of ATRX expression occurs in 20-30% of undifferentiated pleomorphic sarcoma (UPS) and leiomyosarcomas [9, 10]. While the role of ATRX deficiency in the alternative lengthening of telomeres pathway is well described in sarcomas and other ATRX deficient cancers, the chromatin-specific consequences of ATRX loss are not fully understood in the context of disease mechanisms [11, 12].

Sarcomas are mesenchymal neoplasms occurring in connective tissue such as fat, bone, cartilage and muscle [13]. It has been proposed that sarcomagenesis occurs, at least in part, through aberrant differentiation of mesenchymal progenitor cells (MPCs), which has been modeled by introducing sarcoma-relevant driver alterations into MPCs [14-18]. MPCs are multipotent cells that are able to self-renew and undergo differentiation [19] making them well-suited to study perturbations in epigenetic states, which are a key determinants of cell fate and lineage commitment [20-25]. For example, sarcoma-associated mutations in histone genes, which alter histone posttranslational modifications, are sufficient to induce sarcomagenesis when expressed in MPCs [26, 27]. Given the high frequency of *ATRX* alterations in soft tissue sarcoma, we sought to investigate the effect of ATRX deficiency on chromatin and chromatin-dependent processes in the mesenchymal context.

Our findings demonstrate that deletion of ATRX leads to abnormal differentiation in MPCs, which is associated with perturbations in the profiles of histone post-translational modifications and chromatin accessibility in specific regions, which are accompanied by the activation of transposable elements. Our results suggest an important role for the epigenetic regulatory functions of ATRX in the mesenchymal lineage.

## Results

### Loss of ATRX dysregulates transcriptional programs in mesenchymal progenitors

It is well established that the chromatin landscape has a key role in establishing patterns of gene expression, and since ATRX acts as chromatin regulator we hypothesized that ATRX deficiency would lead to changes in transcription. Given that ATRX loss frequently occurs in mesenchymal malignancies, we established *Atrx* KO lines in the C3H/10T1/2 cells, a mesenchymal progenitor cell line, using CRISPR-cas9 with two independent sgRNAs targeting the *Atrx* gene (Fig. S1A). One clone from each guide and an isogenic WT control were selected for further study (Fig. 1A, Fig. S1B). ATRX deletion was confirmed by immunofluorescence and immunoblotting (Fig, 1A, Fig. S1C). Under normal culture conditions without differentiation cues, transcriptomic profiling of *Atrx* KO vs WT cells was performed using bulk RNA-seq. Compared to *Atrx* WT, the *Atrx* KO MPC lines displayed a marked transcriptional dysregulation, with significant gains and losses of gene expression (Fig. S2A, S2B). Focusing on genes that significantly change expression (*p_adj_ <* 0.05) by an absolute magnitude of at least 2-fold in both *Atrx* KO clones when compared to WT, we identified 328 upregulated and 929 downregulated genes (Fig. 1B, 1C, Supplementary Table 1, 2). Given the role of ATRX in establishing and maintaining transcriptionally silenced regions of heterochromatin, we focused on the upregulated gene set. Gene ontology (GO) analysis of the upregulated genes showed an enrichment of gene sets associated with development and differentiation programs, including mesenchymal programs (Fig. 1D, Supplementary Table 3).

**Figure 1:**
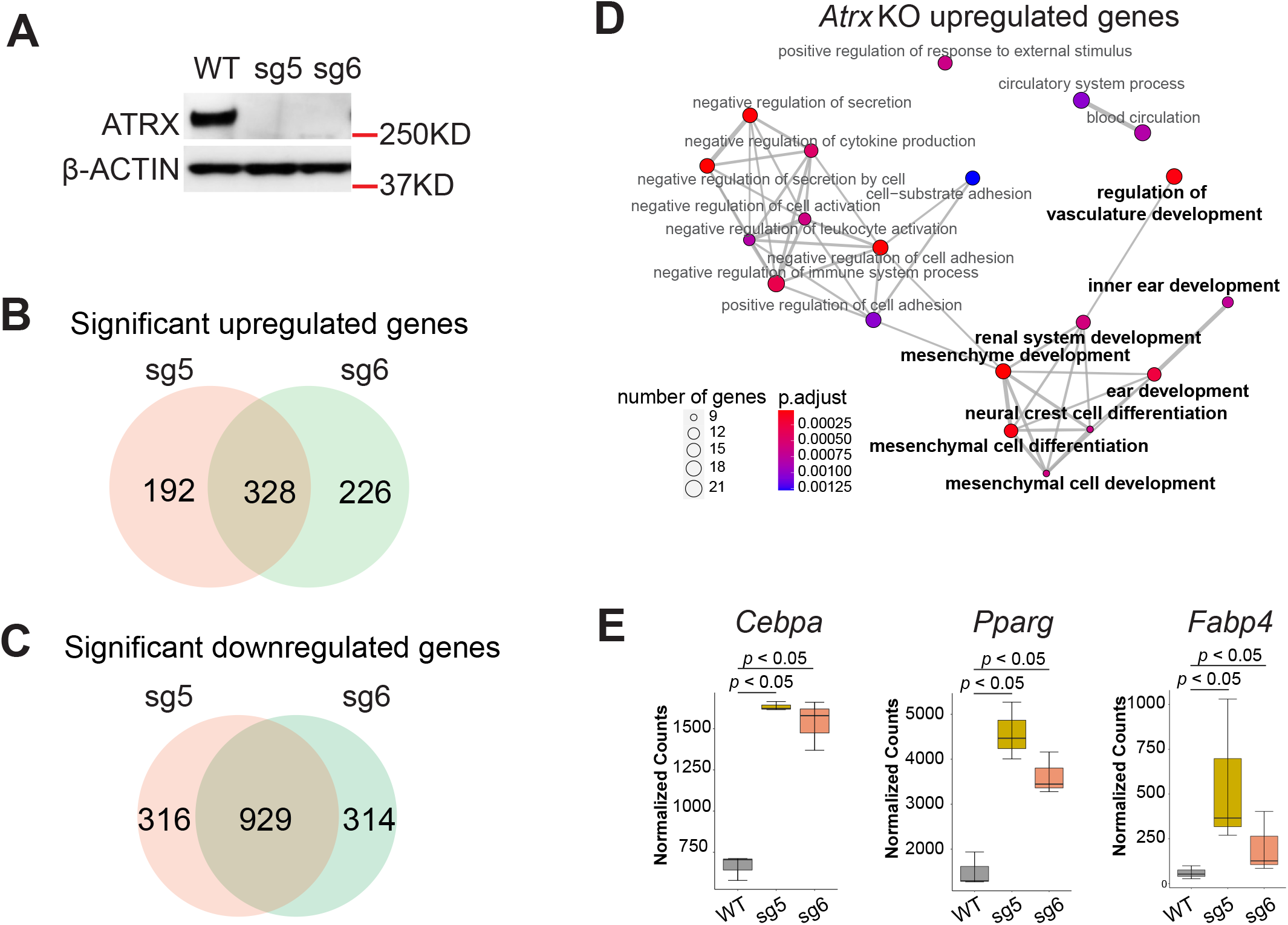
ATRX deficiency alters the transcriptome. **(A)** Immunoblot demonstrating ATRX protein loss in MPC lines. β-ACTIN acts as loading control. **(B)** The intersection of significantly upregulated genes (log_2_foldchange > 1, *p_adj_* < 0.05) in both *Atrx* KO clones based on polyA-RNA-seq datasets. **(C)** The intersection of significantly downregulated genes (log_2_foldchange < -1, *p_adj_* value < 0.05) in both *Atrx* KO clones based on polyA-RNA seq datasets. **(D)** Dot plot of gene ontology (GO) (biological process) analysis for significant upregulated genes from (B). The size of nodes indicates the numbers of genes. The color gradient indicates the *p_adj_* value. Bold type indicates gene ontology terms which are associated with development. **(E)** Boxplot depicting mRNA levels from the RNA-seq dataset. The y-axis indicates RNA-seq read counts normalized by DESeq2. The *p* value (p<0.05) was determined by DESeq2.

To explore the impact of *Atrx* KO on specific mesenchymal lineages, we examined the expression of key adipogenic pathway regulators including the adipogenic transcription factors *Pparγ* and *Cebpa* and the lineage-specific marker *Fabp4*. All three were significantly (log_2_foldchange >1 & *p_adj_* < 0.05) upregulated in *Atrx* KO MPCs (Fig. 1E, Supplementary Table 4), even in the absence of adipogenic differentiation factors. Under the same conditions, we observed significantly decreased expression of mesenchymal stemness markers, *Etv1* [26] and *Cd34* [28], in both *Atrx* KO lines (Fig. S2C). The GO terms associated with downregulated genes were not enriched for pathways relevant for cell lineage differentiation and development (Fig. S2D, Supplementary Table 5).

### ATRX deficiency promotes mesenchymal lineage differentiation and attenuates progenitor properties

The de-repression of developmental and differentiation pathways in *Atrx* KO cells suggests that ATRX may have an important role in restricting differentiation in MPCs. Given the marked upregulation of adipogenic transcription factors and lineage markers in *Atrx* KO lines, we hypothesized that ATRX loss would sensitize MPCs to adipogenic differentiation cues. *Atrx* WT and KO MPCs were treated with adipogenic media to induce differentiation and the degree of differentiation was quantified by staining by Oil Red O (ORO) [29]. Compared to WT MPCs, the *Atrx* KO lines nearly doubled ORO staining intensity after 7 days (p<0.05), indicating that ATRX loss promotes adipogenic differentiation (Fig. 2A). Over the course of the 7-day differentiation treatment, we measured protein expression of PPARγ, C/EBPα, and FABP4 (Fig. 2B). The levels of C/EBP and PPARγ were more highly induced beginning on day 1 of treatment in *Atrx* KO lines compared to WT control whereas the FABP4, a marker for late differentiation, was equally and strongly induced in both settings beginning on day 3. These data suggest that ATRX deletion disturbs the lineage commitment of mesenchymal progenitor cells and promotes differentiation to adipocyte lineage.

**Figure 2:**
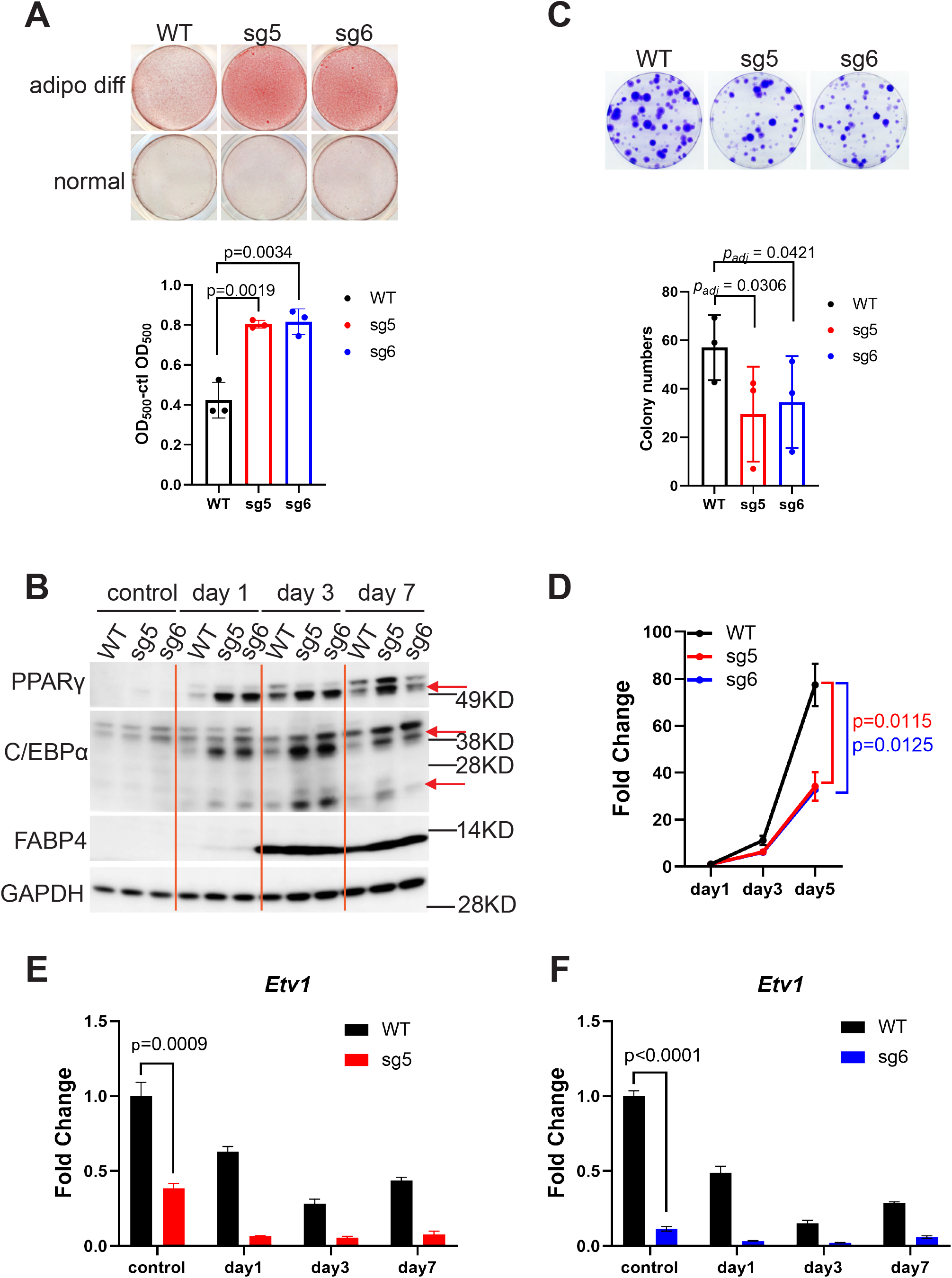
ATRX deficiency promotes differentiation. **(A)** Oil Red O (ORO) staining in adipogenic differentiation experiments. adipo diff: adipogenic media treatment. normal: normal culture media treatment. The y-axis shows the optical density (OD) of the solubilized ORO after adipocyte-specific staining minus the OD value from the normal media control (clt). Data from three biological replicates are plotted with each point representing individual value from each replicate. *p* value was calculated by unpaired *t-test* (two-tailed). **(B)** Western blot of adipocyte markers. The “control” indicates cells grown in the normal media culture at the end of the experiment (Day 7). Day 1, 3, and 7 indicate the duration of adipogenic media treatment. The red arrows indicate the protein bands of interest. Two isoforms of C/EBPα are shown. **(C)** Colony formation assays. The bar plot shows the colony formation from three biological replicates normalized to WT. Cell groups with at least 50 cells were classified as a colony. For each replicate, the percentage of colony formation was normalized to the WT group. The error bars were calculated from three biological replicates. The *p* value was determined by paired one-way ANOVA (WT vs. sg5, WT vs. sg6). **(D)** MPC proliferation assay. Relative numbers of viable cells were determined using an ATPase assay on day 1, day 3 and day 5 after seeding cells. The OD value of luminescence was normalized with day 1 for each group. Three biological replicates were performed. *p* value was calculated by paired *t-test* (two-tailed) for day 5 data. **(E)** and **(F)** RT-qPCR experiments for *Etv1* gene expression in two knockout lines Corresponding to the data in panel (B). *p* value was calculated using one-sample, two-sided *t-test* to compare the mean values for sgRNA samples and WT. The error bars indicate the standard variance.

To assess if *Atrx* KO reduced the stemness properties of mesenchymal progenitors, we performed assays for colony formation (Fig. 2C) and proliferation (Fig. 2D). The two *Atrx* KO lines had reduced colony formation capacity and proliferation compared to *Atrx* WT (p<0.05). We also compared mRNA levels of *Etv1*, a mesenchymal stemness marker, between *Atrx* KO and WT MPCs (Fig. 2E, 2F). The two knock out lines expressed lower levels of *Etv1* during the differentiation time course, including at Day 0, which is consistent with the RNA-seq findings, suggesting that *Etv1* is suppressed at baseline in *Atrx* KO MPCs. These results demonstrate that *Atrx* KO reduces the stem-like properties of mesenchymal progenitors.

### Atrx KO reduces H3K9me3-marked heterochromatin associated with transcriptionally repressed linage commitment genes

Histone post-translational modifications contribute to defining the chromatin states, which in turn impact gene expression [30, 31]. Given that histone post-translational modifications are regulated in part by ATRX, we investigated how ATRX deficiency impacted chromatin states in the MPC model. ATRX, together with DAXX, deposits the histone variant H3.3 at specific genomic regions including telomeres [6, 32, 33]. To confirm loss of ATRX epigenetic function in the knockout cell lines, we performed CUT&RUN (Cleavage Under Targets & Release Using Nuclease) [34] for the histone variant H3.3 in order to compare H3.3 deposition in *Atrx* KO vs WT MPCs. Analysis of three biologic replicates indicated that there were consistent changes in H3.3 localization with gains at some genomic loci and losses at others (Fig. S3A) including significant depletion at telomeres (Fig. S3B), which is consistent with the previous reports [35].

Another known function of ATRX is to establish and maintain locus-specific heterochromatin though recruitment of H3K9 methyltransferases [5]. Using CUT&RUN, we next analyzed how the heterochromatin-associated histone mark, H3K9me3, was altered in *Atrx* KO MPCs. H3K9me3 was significantly changed at specific regions with 4272 gained peaks and 4738 loss peaks (*p* value as 0.05), demonstrating that the ATRX deficiency has effects on heterochromatin in our model (Fig. S3C). Applying GO analysis to genes that are near H3K9me3 differential peaks, we found that many of the top enriched gene sets for regions that had increased or decreased H3K9me3 signal in both *Atrx* KO lines were related to development (Fig. S3D, S3E; Supplementary Table 6, 7).

To further characterize the regions in which H3K9me3 was lost, we first used ChromHMM to annotate chromatin states across the whole genome using CUT&RUN-derived data for H3K4me3, H3K27ac, H3K9me3, and H3K27me3 in *Atrx* WT MPCs (Fig. 3A, S3F, S4) [36]. We found that H3K9me3 was enriched in a state marked by H3K9me3 alone (State 4, Supplementary Table 9), annotated as heterochromatin, as well as in a small subset of regions marked by H3K27me3 and H3K9me3 (State 3, Supplementary Table 8), annotated as repressed chromatin (Fig. 3A). We then compared H3K9me3 CUT&RUN signal within the regions that make up the states containing H3K9me3 in *Atrx* KO and WT MPCs and found that H3K9me3 was reduced in the *Atrx* KO MPCs at these sites (Fig. 3B, 3C). Genes associated with lost H3K9me3 are notably related to cell development and differentiation.

**Figure 3:**
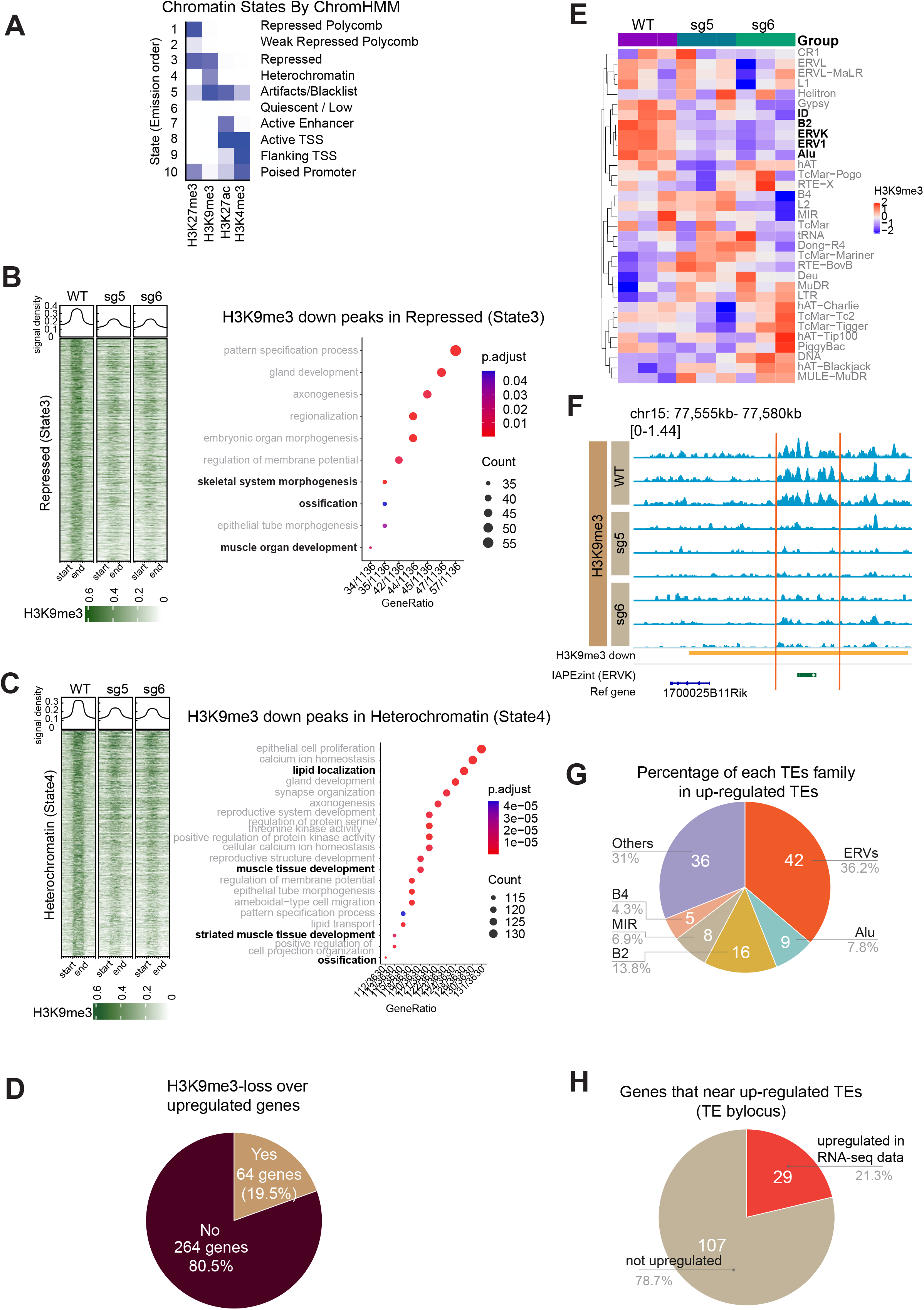
ATRX deficiency reduces H3K9me3 near developmental genes and ERVs in mesenchymal progenitor cells. **(A)** Chromatin states defined by enrichment of histone modifications (H3K4me3, H3K9me3, H3K27ac, and H3K27me3) using ChromHMM. The heatmap shows the emission probabilities of histone modifications in chromatin states. The ten different states. **(B)** The range-based heatmap shows the H3K9me3 signal in repressed regions (State 3) (left) and **(C)** heterochromatin regions (State 4) (left). All peaks are similarly scaled, where “start” and “end” indicate the start and end of the scaled peak. The dot plot shows the GO analysis (biological process) of genes associated with H3K9me3 in each chromatin state (right). **(D)** Percentage of H3K9me3 lost peaks (*p*<0.05) associated genes (closest gene to peak) that do (“yes”) or do not (“no”) overlap with up-regulated genes (log_2_foldchange > 1 & *p_adj_* < 0.05). **(E)** Heatmap of H3K9me3 signal at repetitive regions in *Atrx* WT vs KO MPCs derived by SQuIRE analysis of three biologic replicates. Bold indicates elements of interest (see main text). **(F)** IGV tracks show a representative H3K9me3 differential peak at an IAPEzint ERV element. The yellow bar indicates a differential H3K9me3 peak. The green bar shows the mouse IAPEz-int element. Within each sample genotype, each track represents an independent biologic replicate. **(G)** Percentages of each family of TE up-regulated in *Atrx* KO vs WT MPCs based on an rRNA-depletion sequencing dataset. The up-regulated transposable elements were mapped at an individual locus level. **(H)** Percentage of up-regulated genes in polyA-RNA-seq data that annotated by up-regulated TEs (TE method by locus).

**Figure 4:**
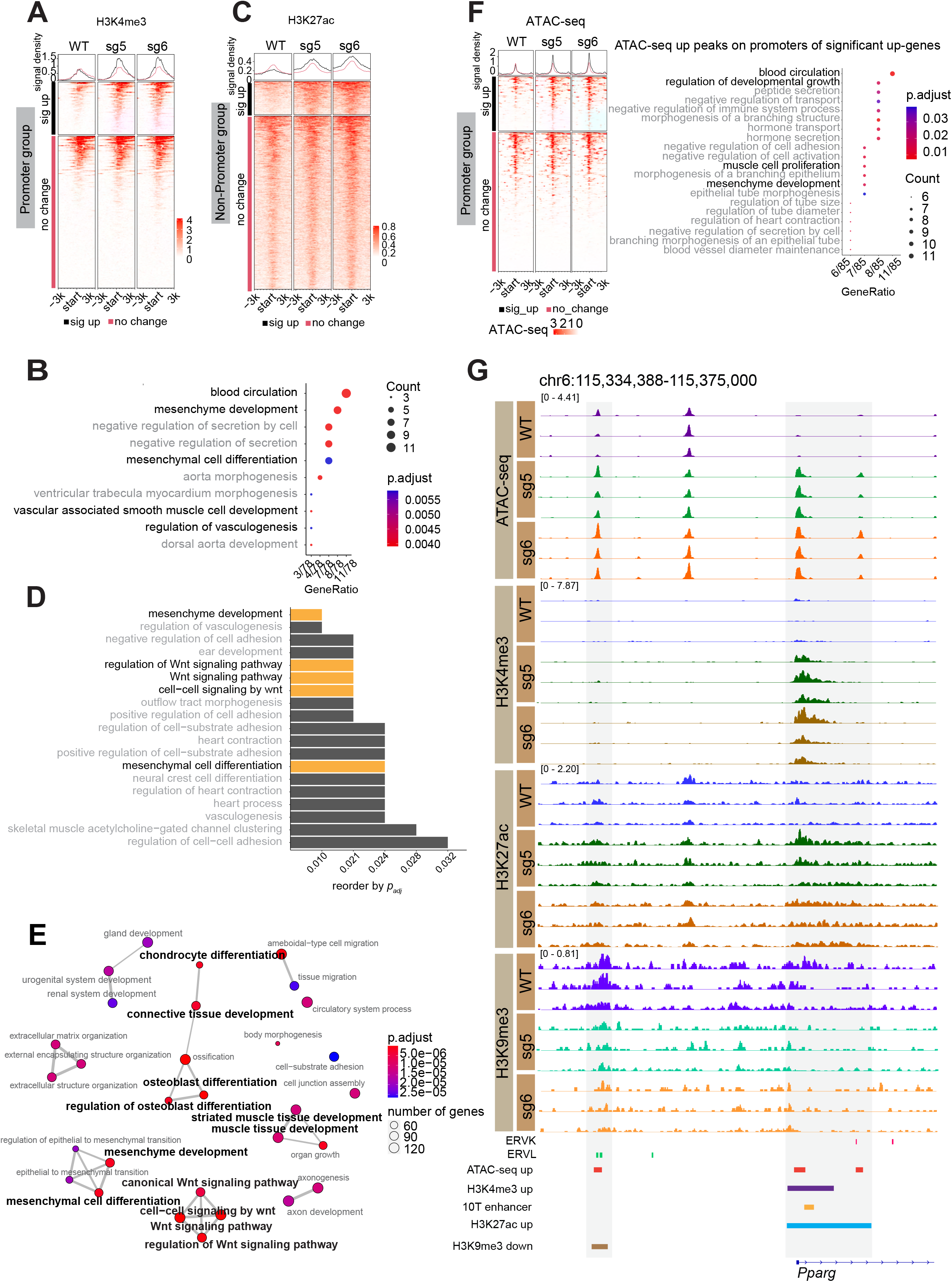
Loss of ATRX alters active chromatin marks and perturbs chromatin accessibility to induce expression of a key adipogenic regulator. **(A)** H3K4me3 signal around promoters of significantly upregulated genes grouped by whether H3K4me3 is gained (black bar) or unchanged (red bar) **(B)** GO terms (biological process) associated with H3K4me3 gained regions. **(C)** H3K27ac signal +/- 3 kb around the center of H3K27ac peaks that do not overlap promoters grouped by whether H3K27ac is gained (black bar) or unchanged (red bar) **(D)** GO analysis (biological process) of H3K27ac significantly increased regions. **(E)** Network plot of gene programs associated with increased accessibility in *Atrx* KO lines. The terms labeled with bold black font indicate those related to the mesenchymal lineage. **(F)** Signal of ATAC-seq +/-3 kb around the transcription start site (TSS) of genes that have significantly increased expression in *Atrx* KO MPCs and have an ATAC-seq peak in the TSS region. The regions are grouped based on whether the overlapping ATAC peak has significantly increased signal (black bar) or is unchanged (red bar). The dot plot shows the GO terms associated with upregulated genes. **(G)** IGV tracks of CUT&RUN and ATAC-seq replicates showing a region near the promoter of the *Pparγ* gene. Enhancer annotation is from EnhancerAtlas 2.0 database (http://www.enhanceratlas.org/) [52] and converted to the mouse mm10 genome.

Next, to better understand the consequences of this observation on gene regulation, we examined the connection between H3K9me3 loss and changes in gene expression. Of the 328 significantly upregulated genes in *Atrx* KO MPCs, 19.5% (64/328) were associated with peaks that had significantly less H3K9me3 signal in the ATRX deficient line (Fig. 3D). GO pathway analysis of these genes revealed enrichment of gene sets related to adipocyte differentiation, including lipid localization, fat cell differentiation, and brown fat cell differentiation (Supplementary Table 10). These genes included key adipogenic regulators and markers such as *Fabp4* and *Pparγ*, which are part of the fat cell differentiation GO gene set (Supplementary Table 10). In addition, another gene in the same gene set, *Lpl*, has roles in adipogenic metabolism by mediating hydrolysis of circulating lipoprotein particles [37]. Examination of integrated genomic viewer (IGV) tracks in a putative *Fabp4* regulatory element demonstrates H3K9me3 depletion in *Atrx* KO cells (Fig. S3G).

These observations indicate that ATRX deficiency reduced the heterochromatic histone mark H3K9me3 at genomic regions that were associated with genes relevant to linage-specific differentiation and development. These data suggest that H3K9me3 loss may contribute to gene expression changes in ATRX deficient MPCs, leading to their enhanced capacity for adipogenic differentiation.

### Atrx KO reduces H3K9me3 at ERV repetitive regions

ATRX and the ATRX-associated H3K9 methyltransferase, SETDB1, promote H3K9me3 deposition and silencing at repetitive elements [5]. These sequences include transposable elements (TEs) such as endogenous retroviral elements (ERVs), which are a type of long terminal repeat derived from integrated retroviral elements [38]. De-repression of ERVs can lead to formation of dsRNA, which is detected by sensors that in turn stimulate innate immune signaling, which has been implicated as a mediator of antitumor immune response [39-42]. Separately, ERVs can act as gene regulatory elements with enhancer-like features and as transcription factors binding sites to regulate gene expression [43-45]. Given that H3K9me3 is depleted in heterochromatin regions in *Atrx* KO MPCs, we sought to determine how this affected ERV expression and gene regulation in this system.

We compared H3K9me3 enrichment at repetitive elements between *Atrx* KO and WT MPCs. We found that H3K9me3 signal was reduced by deletion of ATRX on multiple types of TEs, including ERV family members (ERV1, ERVK), *Alu*, *ID* and *B2* (Fig. 3E). To further define the relationship between H3K9me3 loss at repetitive elements and differentiation phenotypes in *Atrx* KO lines, we mapped the repetitive elements onto the previously defined chromatin states (Fig. 3A). While most of repetitive elements were in quiescent (State 6) regions devoid of detectable chromatin marks (Fig. S5A), outside of these regions ERV1 and ERVK elements preferentially mapped to heterochromatin regions where H3K9me3 was reduced in *Atrx* KO vs WT MPCs (State 4) (Fig. S5B).

To identify specific subfamilies of ERVs where H3K9me3 is depleted in *Atrx* KO MPCs, we analyzed the H3K9me3 differential peaks by adapting SQuIRE analysis, a RNA-seq analysis pipeline that provides a quantitative and locus-specific information on TE expression [46]. The top reduced regions of H3K9me3 in both *Atrx* KO lines is an IAP element (Fig. S6A, S6B, Supplementary Table 11,12), which is a member of the ERVK family. Loss of H3K9me3 in *Atrx* KO vs. WT at a representative region containing an ERVK (IAPEzint) element can be appreciated by review of IGV tracks (Fig. 3F).

To determine if TEs are transcriptionally de-repressed in the setting of ATRX-dependent H3K9me3 reduction, we performed RNA-sequencing following rRNA-depletion and analyzed locus-specific repetitive element differential expression between *Atrx* KO and WT MPCs using SQuIRE. We identified 116 significant (*p_adj_* <0.05) upregulated TEs in common between both *Atrx* KO clones (Supplementary Table 13). Among these upregulated TEs, 42 (36.2%) belong to the ERV superfamily (Fig. 3G), which is the largest subset of upregulated TEs.

To investigate whether de-repressed TEs correlated with nearby gene expression, we analyzed the intersection of upregulated TEs with significantly upregulated genes. We found that the 116 significant upregulated TEs were annotated by 136 genes (using the TE transcript by locus method, Supplementary Table 14). Among those genes, 29 genes (21.3%) were significantly upregulated as measured by RNA-seq (Fig. 3H). The adipocytic lineage transcription factor, *Pparγ*, was observed in this gene set, raising the possibility that the associated TE could serve as a regulatory element for *Pparγ*. In addition, we observed 445 significantly downregulated TEs near 163 genes, of which 30 genes were also downregulated (Fig. S6C, Supplementary Table 15). These results demonstrate that *Atrx* KO leads to reduction of H3K9me3 on specific TEs including those in regions, which potentially regulate an adipogenesis master regulator.

### Loss of ATRX promotes chromatin accessibility and enrichment of active histone modifications at developmental genes

ATRX regulates euchromatin as well as heterochromatin [47], leading us to investigate whether the dysregulation of developmental transcriptional programs in MPCs that we see in *Atrx* KO was connected to changes in active histone modifications, such as H3K4me3 and H3K27ac. We hypothesized that loss of ATRX leads to gain of H3K4me3 in specific promoters and H3K27ac in enhancer-like regions. First, we investigated the distribution of H3K4me3 and H3K27ac in *Atrx* WT and KO MPCs and identified genomic features of differential peaks of H3K4me3 and H3K27ac in *Atrx* KO. As expected, H3K4me3 gained peaks in *Atrx* KO clones are highly represented at promoters (Fig. S7A, Supplementary Table 16). Enrichment of H3K27ac was mainly in introns and distal intergenic regions in *Atrx* KO lines (Fig. S7B, S7C, Supplementary Table 17).

Next, we examined the relationship between distribution of H3K4me3 at promoter regions and transcriptional profiles of those genes. Interestingly, less than half of genes upregulated in *Atrx* KO cells gain H3K4me3 at their promoters (Fig. 4A) suggesting that other factors may mediate the increased expression in the group where H3K4me3 does not change. Focusing specifically on genes that gain H3K4me3 and are upregulated in an ATRX-dependent fashion, GO pathway analysis revealed genes related to mesenchymal differentiation and which included the adipogenesis master regulator, *Pparγ* (Fig. 4B, Supplementary Table 18).

We next examined the distribution of H3K27ac, focusing on non-promoter regions near upregulated genes since in these regions, H3K27ac is commonly attributed to active enhancers. Similar to our findings with H3K4me3 in ATRX deficient cells, H3K27ac gain was seen in a minor subset of upregulated genes (Fig. 4C). However, these genes again were enriched in GO terms related to mesenchymal development, including Wnt signaling, which is important for mesenchymal differentiation [48-50] (Fig. 4D, Supplementary Table 19). Thus, aberrant enhancer activation following ATRX loss could contribute to the pro-differentiation phenotype in *Atrx* KO MPCs.

Our data suggests that *Atrx* KO in MPCs leads to changes in both the heterochromatic histone mark H3K9me3 and the euchromatic marks H3K4me3 and H43K27ac. Given the connection of these posttranslational modifications with chromatin compaction, we speculated that ATRX deficiency would lead to a shift in chromatin accessibility. To test this, we performed ATAC-seq, which demonstrated increased accessibility in *Atrx* KO cells at genes associated with mesenchymal development and differentiation (Fig. 4E, Supplementary Table 20). To investigate the connection of increased chromatin accessibility driven by ATRX deficiency and gene expression changes in *Atrx* KO clones, we separated ATAC-seq peaks at promoters of significantly upregulated genes (Supplementary Table 21) from those at non-promoter regions near or in the same genes (Supplementary Table 22) (Fig. 4F). Similar to H3K4me3, while a minority of the upregulated genes showed increased accessibility at their promoters (Fig. 4F), these genes were associated with development. The observation that a group of genes increased expression but did not gain chromatin accessibility suggests, as is in the analogous case of upregulated genes that did not gain H3K4me3, that non-chromatin mechanisms may account for transcriptional changes upon deletion of ATRX.

To understand the relationship between heterochromatin changes and increased chromatin accessibility in ATRX deficient MPCs, we intersected the ATAC-seq gained peaks (8742 peaks) that overlapped with regions showing decreased H3K9me3 (4568 peaks) and identified 170 overlapping regions (Fig. S8A). Despite this small number of overlapping peaks, GO analysis suggested that this subset was highly enriched for programs related to development including in along the mesenchymal lineage (Fig. S8B, Supplementary Table 23). These results demonstrate that the increased chromatin accessibility coupled with a reduction in H3K9me3 at specific genes in *Atrx* KO MPCs is associated with altered gene expression and a pro-differentiation phenotype in *Atrx* KO MPCs.

### ATRX deficiency induces an active chromatin state at the promoter and putative regulatory element of the adipogenic transcription factor Ppar

ATRX loss leads to an aberrant increase in expression of *Pparγ* (Fig. 1E), which is an important adipogenic transcription factor [51]. In addition, in functional assays for adipocytic differentiation, PPARγ protein levels are markedly induced in *Atrx* KO MPCs immediately after induction of adipogenic differentiation (Fig. 2B). In order to understand how these phenotypes are functionally linked to the epigenetic changes driven by ATRX deficiency, we investigated ATRX-dependent changes in the chromatin state near the *Pparγ* gene. In *Atrx* KO MPCs the promoter of *Pparγ* showed increased accessibility and increased H3K4me3 and H3K27ac, suggesting a more active chromatin state (Fig. 4G). Notably, previous work has identified a *Pparγ* enhancer element in this same region [52] (Fig. 4G, Supplementary Table 24), suggesting multiple mechanisms by which these chromatin changes could enhance *Pparγ* expression. In addition, we observed a loss of H3K9me3 and increased accessibility at an ERVL element 20 kb upstream from the *Pparγ* promoter, raising the possibility that it functions as an ATRX-dependent gene regulatory element for *Pparγ*.

### ATRX associates with accessible and active chromatin in MPCs

To understand whether direct ATRX binding could influence the gene expression and chromatin changes observed in *Atrx* KO MPCs, we mapped ATRX binding sites using CUT&RUN. At telomeric regions, where ATRX is known to function with DAXX to deposit H3.3 and bind to G-quadruplex structures [6], ATRX signal was decreased in both knockout clones compared to WT (p < 0.05) (Fig. S9A). As an additional control, we detected ATRX peaks at zinc finger gene clusters (Fig. S9B), where ATRX is known to bind [53] that were lost in the *ATRX* KO MPC.

ATRX enrichment at the sites of active transcription start sites (TSS) was recently reported in lymphoblastoid cell lines, suggesting that ATRX has an important role in euchromatin where it regulates gene expression in addition to its more established role at heterochromatin [47]. This is consistent with MPC ATRX peaks identified here (n=114; *p_adj_* < 0.05) (Fig S9C, Supplementary Table 25, 26), where we observed that of the 114 ATRX binding sites, 33.3% were located at promoter regions, and 38.6% were in distal intergenic regions (Fig. 5A). The remainder associated with 5’UTRs (5.3%), exons (7.9%), 3’UTRs (2.6%), and introns (12.3%). Mapping ATRX binding sites onto the ChromHMM-defined chromatin states revealed an association with active transcription start sites (TSS, State 8) (Fig. 5B). Comparing the ATRX binding sites to *Atrx* WT ATAC-seq peaks, we found that 87.7% (100/114) of ATRX binding sites overlapped with chromatin accessible regions (Fig. 5C, Supplementary Table 27). ATRX binding corresponded to H3K4me3 enriched regions in 70.1% of sites (80/114) (Fig. 5D, Supplementary Table 28) and with H3K27ac in 81.5% (93/114) (Fig. 5E, Supplementary Table 29). Interestingly, in the intersection of H3K27ac occupied regions with ATRX binding sites, 66.7% (62/93) were on non-promoter regions, suggesting a potential role for a subset of ATRX at active enhancers in MPCs (Fig. S9D, Supplementary Table 30).

**Figure 5:**
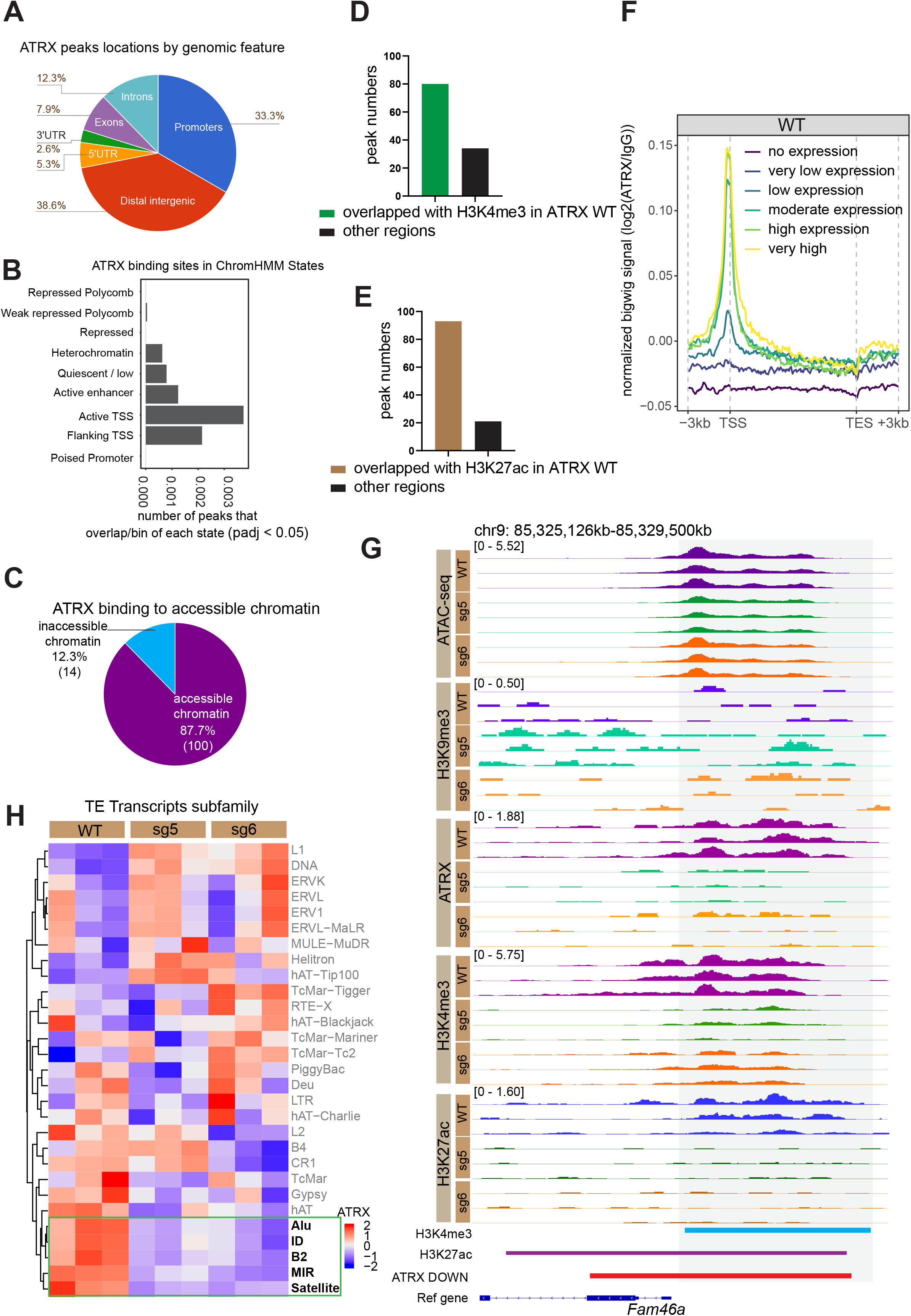
ATRX binding to active chromatin. **(A)** ATRX binding sites by genomic feature. **(B)** Overlap of ATRX binding sites with chromatin states excluding state 5 (artifact/blacklist). x-axis indicates the level of ATRX binding calculated by: (number of ATRX peaks that overlap state bins)/(total bins with that state) **(C)** The percentage of ATRX peaks that overlap ATAC-seq peaks. **(D)** The numbers of ATRX peaks that overlap H3K4me3 peaks in *Atrx* WT MPCs. **(E)** The numbers of ATRX peaks that overlap H3K27ac peaks in *Atrx* WT MPCs **(F)** ATRX signal near TSS in gene groups binned based on expression levels. The genes were grouped as follows: “no expression” indicates that TPM (Transcripts Per Million) = 0, “very low expression” indicates the quintile of TPM between 0-20%, “low expression” indicates the quintile of TPM between 20-40%, “moderate expression” indicates the quintile of TPM between 40-60%, “high expression” indicates the quintile of TPM between 60-80%, and “very high expression” indicates the quintile of TPM between 80-100%. **(G)** IGV tracks at the *Fam46a* gene region. **(H)** Heatmap of ATRX signal over repetitive elements.

Given that ATRX peaks were often localized to open chromatin and active promoters, we explored the relationship between ATRX binding and gene expression. Genes were binned based on expression and ATRX signal was quantified around the TSS of genes in each group (Fig. 5F). The most highly expressed genes had the greatest ATRX signal, whereas non-expressed genes had no ATRX enrichment at the TSS. For example, *Fam46a*, which is known to downregulated in differentiating adipocytes [54] and was significantly downregulated in *Atrx* KO MPCs (Fig. S9E) has an ATRX binding site at its promoter region (Fig. 5G). This association of ATRX with active genes is similar to a recent report in a hematopoietic lineage [47] and suggests that ATRX may play a role in maintaining and potentially establishing gene expression in the mesenchymal cells. However, given that only a subset of ATRX binding sites correlate with changes in gene expression in *Atrx* KO MPCs (Supplementary Table 28), there are likely other mechanisms by which ATRX regulates transcription. One possibility may be an indirect mechanism via binding and chromatin-mediated regulation of transposable elements which in turn influence transcription. In support of this possibility, we observed that ATRX binds several families of repetitive elements, including SINEs (*Alu*, *ID*, B2, MIR) and Satellite DNAs (Fig. 5H), a finding that was confirmed by a second analysis method (Fig. S9F).

## Discussion

The mesenchymal lineage gives rise to connective tissues. Sarcomas, which are cancers of connective tissues, have recurrent loss of ATRX in up to 30% of specific subtypes, suggesting the potential importance of ATRX deficiency in sarcoma biology and possibilities for precision medicine approaches to treatment [7]. However, while ATRX loss is known to contribute to the alternative lengthening of telomere phenotype in sarcomas, the functions of ATRX in the epigenetic regulation of gene expression and downstream processes in the mesenchymal context is less understood.

Given the important role for ATRX regulating epigenetic states and the intersection of epigenetics with development, we hypothesized that loss of ATRX would affect cell commitment in mesenchymal progenitor cells. We demonstrated that the *Atrx* KO MPCs were sensitive to differentiation induction and exhibited reduced progenitor properties. At the transcriptional level, ATRX loss increased expression of adipogenesis regulators and markers, including *Pparγ*, which is an early and direct master regulator of adipogenesis [51]. We then investigated ATRX mediation of these phenotypes through epigenetic changes, with a focus on several potential and non-exclusive mechanisms (Fig. 6).

**Figure 6:**
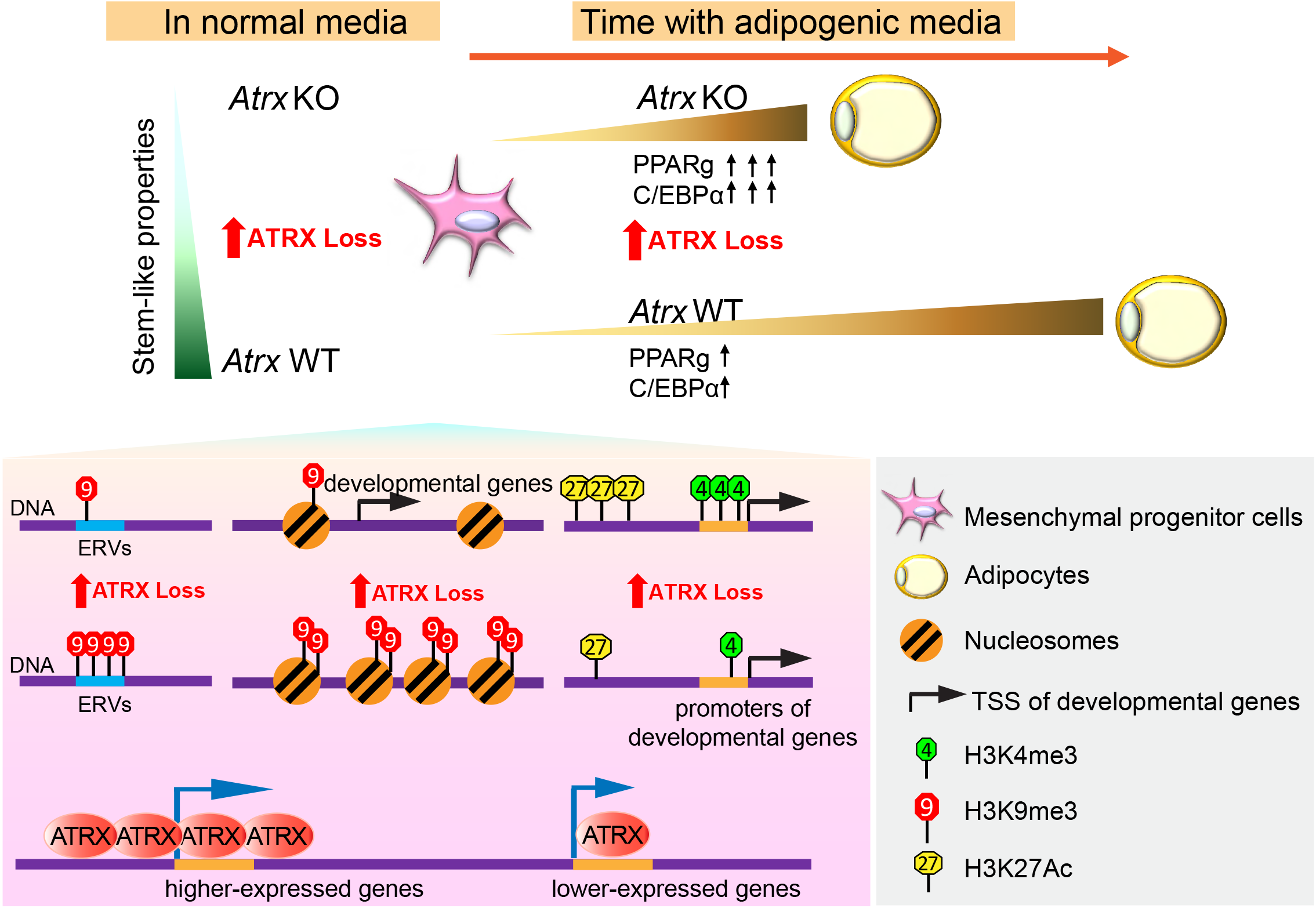
Model for ATRX-dependent chromatin and gene regulation in MPCs. *Atrx* KO MPCs demonstrate reduced stem-like properties, such as slower proliferation, reduced colony formation and downregulated mesenchymal stemness gene expression. In adipogenic media, *Atrx* KO MPCs exhibit accelerated adipocytic differentiation and expression of adipogenic transcription factors. We propose a potential model in which these phenotypes are regulated by multiple epigenetic mechanisms at developmental genes including loss of H3K9me3 near promoters and at ERVs with putative cis-regulatory functions, gain of chromatin accessibility, and enrichment of active histone marks. In addition, loss of direct binding of ATRX, which is associated with high levels of genes expression, may contribute. Together, these mechanisms could explain gene expression changes in ATRX deficient MPCs, which in turn mediate cell differentiation in the mesenchymal lineage context.

First, ATRX deficiency promotes chromatin accessibility and increased transcription at genes associated with mesenchymal differentiation including *Pparγ* and *Fabp4*. Whether ATRX regulates chromatin accessibility directly or indirectly remains an interesting question to be addressed in future work. Second, the loss of ATRX leads to changes in histone posttranslational modifications at specific loci, including the heterochromatin mark H3K9me3 and active chromatin marks, H3K4me3 and H3K27ac. The genes associated with H3K9me3-depleted heterochromatin regions in ATRX-deficient cells are relevant for differentiation and development. This suggests that loss of this repressive mark may create a chromatin environment permissive to signals that activate these programs, which is consistent with the accelerated and increased differentiation phenotype observed in *Atrx* KO MPCs following simulation with adipogenic media. In addition, ATRX deficiency leads to the loss of H3K9me3 at ERVs, which could represent another potential mechanism for the transcriptional changes induced by ATRX loss given that ERVs can serve as cis-regulatory elements. Finally, histone marks associated with active transcription were also altered upon ATRX loss. We observed increased H3K4me3 at promoters and H3K27ac at putative enhancer (i.e., non-promoter) regions near significantly upregulated genes. This suggests that increased H3K4me3 and H3K27ac may also contribute to the induction of mesenchymal development programs, such as Wnt-associated signaling [48-50].

Third, ATRX may regulate gene expression through direct interactions with active chromatin. Recent reports in lymphoblastoid cell lines shows that 38% of ATRX binding sites are localized to promoters [47], which is consistent with our results demonstrating that approximately one-third of ATRX binding sites in MPCs occur at promoters. In contrast, ATRX binding sites in murine embryonic stem cells are mainly restricted to distal intergenic regions and gene bodies, with only 1% on promoters [55]. While in murine neuroepithelial progenitors, 17% of ATRX binding sites were located in promoters [55]. Therefore, ATRX binding sites differ depending on the cell and developmental contexts with a trend towards increased association with promoters in lineage specific progenitors compared to less differentiated stem cells. Our findings demonstrate that ATRX is not only required for maintaining heterochromatin, but that it may also regulate gene expression through binding to active chromatin regions, consistent with this recently discovered role for ATRX. In our system, ATRX binding was associated with increased gene expression. However, whether ATRX occupancy directly drives transcriptional upregulation is unclear since only a subset of genes near ATRX binding sites are downregulated following ATRX loss. We speculate that ATRX binding might regulate gene expression changes through a distal enhancer mechanism in the case of ATRX binding sites that do not influence expression of nearby genes. In support of this, ATRX signal was decreased at several types of SINEs in *Atrx* KO MPCs, including *Alu*, *B2*, *B4*, *ID*, and MIR, which act as transcription factor binding sites (*Alu*[56], MIR[57]) or harbor promoter-like (*B2*) [58] or enhancer-like features (*B4)* [59], respectively.

Our findings demonstrate that ATRX restricts differentiation in mesenchymal progenitor cells. Deletion of ATRX leads to aberrant de-repression of the lineage-restricted transcriptome, accompanied by changes in chromatin accessibility and histone post-translation modifications near transposable elements and at specific genes associated with differentiation. These findings raise several questions including whether the changes observed in active histone marks and chromatin accessibility after *Atrx* KO is a direct or indirect effect and how these findings are relevant to mesenchymal malignancies where ATRX loss is common (e.g., undifferentiated pleomorphic sarcoma and leiomyosarcoma). One intriguing possibility is that ATRX loss creates a permissive chromatin and transcriptional context in which mesenchymal progenitors are susceptible to subsequent sarcomagenic events such as loss of *TP53* and *RB1* [7]. Overall, our results expand our understanding of ATRX function in in the mesenchymal context.

## Acknowledgments

We are very grateful to Prof. C. David Allis (CDA), who unfortunately passed away in January 2023, for his support, invaluable suggestions, and insight. We also thank Allis lab members for suggestions and advice. We also acknowledge Samer Shalaby from the Flow Cytometry Resource Center at the Rockefeller University for the cell sorting assistance. The CUT&RUN sequencing and Omni-ATAC sequencing were performed by Rockefeller University genomic core. This work was supported by a Connective Tissue Oncology Society Basic Science Research Award (BAN, CDA, WDT), NCI K08 CA245212 (BAN), the Memorial Sloan Kettering Cancer Center Support Grant (P30 CA008748), and the University of Pittsburgh Hillman Cancer Center Support Grant (P30CA047904).

## Methods

### Cell culture

C3H/10T1/2 (CCL-226) cells (MPCs) were obtained from ATCC. Except where otherwise stated for specific experiments, cells were cultured in DMEM (Corning, 10-013-CV) plus 10% FBS (ATLANTA biologicals, S11150, heat inactive) with 1% penicillin/streptomycin (Gibco, 15140-122) at 37-degree Celsius culture condition with 5% CO_2_. The parental cell line was tested for mycoplasma and authenticated by ATCC.

### CRISPR-Cas9 cloning

The vector for CRISPR-cas9 pSpCas9(BB)-2A-GFP (PX458) (Addgene, #48138) was obtained from Addgene. Mouse *Atrx* sgRNAs were designed using Benchling (https://www.benchling.com/). Target sequences: sg5: 5’ TGGCCGTAAAAGTTCTGGGG-3’, sg6: 5’-CTACTGGACTTGGTGACTGC-3’. The control group was generated using an empty vector without sgRNA but carrying Cas9. The vector was digested by Bbsl (NEB, R0539S) and dephosphorylated by calf intestinal alkaline phosphatase (NEB, M0290). The oligos were annealed with T4 ligation buffer (NEB, B0202S) with T4 PNK (NEB, M0201L) enzyme. The ligation step was performed using Quickligation buffer (NEB, B2200S) and Quick Ligase (NEB, M2200L). The ligation products were expressed in Stbl3 competent cells (Thermofisher, C737303).

### Transient transfection

Two micrograms of plasmid products carrying sgRNAs with Cas9 were transfected using Lipo2000 (Thermofisher, 11668019) into 0.7 × 10^4^ MPCs cells. The transfection procedure was according to the manufacturer’s instructions. Cells were harvested 48 h after transfection for sorting and sorted into 96 well plates using BD FACSAria II Cell Sorter based on GFP signal.

### Proliferation assay

50 *Atrx* WT and *Atrx* KO cells were grown in 96-well plates (Costar, 3917) for 5 days. Proliferation was evaluated using the CellTiter-Glo Luminescent Cell Viability Assay kit (Promega, G7572), following the manufacturer’s instruction. The luminescence was read immediately in 96-well plate reader (BioTek, Synergy hybrid H4 Reader).

### Colony formation

100 C3H/10T1/2 cells (WT and KO clones) were seeded in 10 cm dishes, cultured in α-MEM (Corning, 10-009-CV) with 20% FBS (ATLANTA biologicals, S11150), with 1% penicillin/streptomycin (Gibco, 15140-122). Cultures were incubated at 37-degree Celsius with 5% CO_2_ for 10–14 days. When visible colonies formed, colonies were washed once with PBS. Cells were stained with 0.5% crystal violet (SIGMA-ALDRICH, C3886) in 4% formaldehyde (Fisher chemical, F79-500) for 1 h at RT. After removing the staining buffer, cells were washed with ddH_2_O for 3-5 minutes to remove the background staining and then digitally imaged (Epson Perfection V800 photo). Colonies were manually counted. A group of cells had to contain at least 50 cells to be considered a colony.

### Adipogenic differentiation

C3H10T1/2 were differentiated into mature adipocytes following treatment with insulin, dexamethasone, troglitazone, and methylisobutylxanthine per an established protocol [29]. 1 × 10^5^ C3H/10T1/2 were seeded in 24-well plates such they were fully confluent. Adipocyte differentiation medium [29] consisting of DMEM (Corning, 10-013-CV) with 10% heat inactive FBS (ATLANTA biologicals, S11150), 0.5 mM isobutyl methylxanthine (Sigma, I7018), 1 µM dexamethasone (sigma, D4902), 5 µg/mL insulin (Sigma, I9278), 5 µM troglitazone (Sigma, T2573) was used to replace the media the following day. After 2 days, the media was changed every 2 days with insulin and FBS the only additive to the DMEM. Differentiation was assessed by Oil Red O after day 7.

### Oil Red O staining

The Oil Red O staining was performed as previously described [29]. 0.5% w/v Oil Red O stock (Sigma, O0625) in 100% isopropanol (Fisher chemical, A416SK-4) was prepared fresh. A working solution was prepared by mixing contained 6 mL of stock solution and 4 mL of H_2_O, followed by filtering (0.45 µM, Pall Corporation syringe filter, PN-4614) to remove precipitate. Cells were washed by PBS. Cells were fixed with 4% formaldehyde in PBS for 2 minutes at room temperature, followed by washing with H_2_O. Next, 220 µL Oil Red O working solution was added to the wells and incubated cells for 1 h at room temperature. Then cells were washed twice with 0.5% isopropanol to remove background staining. Plates were digitally imaged (Epson Perfection V800 photo) before the Oil Red O was solubilized by adding 650 µL of 100% isopropanol to the stained cells, which were incubated for 20 minutes on a shaker. The eluate was transferred into a 96 well plate (Corning, 3361) and the absorbance measured at 500 nm with 96-well plate reader (BioTek, Synergy hybrid H4 Reader).

### Immunofluorescence

800 C3H/10T1/2 cells were seeded in a 96 well plate (glass bottom culture plates, MatTek, PBK96G-1.5-5-F) and keep cells in 37-degree for 24 h. Cells were fixed with 4% formaldehyde in PBS for 10 minutes at room temperature. Cells were permeabilized with 0.2% digitonin (EMD Millipore, 300410) in PBS (Corning, 21-040-CV) for 10 minutes at room temperature. Next, the cells were incubated in 3% BSA (in PBS, Sigma, #9418) for blocking for 1 h at room temperature followed by incubation in primary antibody (ATRX, Santa Cruz, sc-15408, 1:200) at 4-degree overnight, followed by secondary antibody (1:600, Invitrogen, A32754) incubation at room temperature for 1 h. Cells were washed three times with PBS for 5 minutes followed by DAPI staining (2 µg/ml in PBS, Sigma, D9564) for 5 minutes at room temperature. Mounting media (Vectashield, H-1000) was added immediately after DAPI staining. Cells were imaged using Widefield Microscope CellDiscoverer7 (CD7) automated widefield high-throughput system (Zeiss). Images were processed with ImageJ software (http://rsb.info.nih.gov/ij/). For the ATRX antibody, the ImageJ Brightness/Contrast was set as 30/112. For DAPI, the Brightness/Contrast was set as 37/123. All images were processed with the same parameter settings for each antibody.

### Protein isolation and Western Blot

Cell pellets were lysed in lysis buffer NETN (20 mM Tris (pH 7.5), 1 mM EDTA, 150 mM NaCl, 0.5% NP-40, protease inhibitor tablet (Roche), 0.5 mM DTT). Samples were incubated in the cold room for 30 minutes followed centrifugation at 4-degree Celsius with maximum speed for 10 minutes. The supernatant was collected and the concentration determined by BCA quantification (Pierce BCA protein Assay kit, Cat.23225, Thermo scientific) to allow normalization between samples for Western blotting. Samples were mixed with Laemmli Sample Buffer (4X) (containing 1.0 M Tris-pH6.8, 8% SDS, Glycerol, β- Mercaptoethanol (10%), Bromophenol blue) for boiling at 100-degree Celsius for 10 minutes. The samples were then separated by SDS-PAGE and analyzed by standard immunoblotting using running buffer from Invitrogen (NuPAGE™ Tris-Acetate SDS Running Buffer (20X), LA0041) and transfer buffer from Thermofisher (NUPAGE transfer buffer, NP00061). The blotting processes was performed as previous described [26]. Antibodies used for Western blot are the following: ATRX (Santa Cruz, sc-15408, 1:800), FABP4 (R&D, AF1443, 1:5000), C/EBPα (CST, #2295, 1:1000), PPARγ (CST, #2430), β-actin (Abcam, ab8224), GAPDH (Abcam, ab8245, 1:1000).

### Reverse Transcriptase Quantitative PCR (RT-qPCR)

For RT-qPCR, RNA was prepared with RNeasy Mini kits (Qiagen, 74104). The RNAs concentration was determined using a Nanodrop (Spectrophotometer, ND-1000). cDNA was prepared published procedures [26]. qPCRs were performed using SYBR green PCR master mix (Applied Biosystems, 4367659). The detailed steps are following previous described [26]. The endogenous control gene was 18S. The sequences of primers are as following: mouse-Etv1: F:5’-GTTTGTTCCAGACTATCAGGCTG-3’, R: 5’-GGGCTGTGGGGTTCTTTCTT-3’. mouse-18s: F:5’-GTAACCCGTTGAACCCCATT-3’, R: 5’-CCATCCAATCGGTAGTAGCG-3’. The statistical analysis was performed using a one-sample, two-sided *t-test*.

### PolyA-RNA-seq and data analysis

Approximately 1 million cells were collected from 10 cm dishes. RNA extraction, polyA selection, library preparation, and RNA sequencing were performed at the MSKCC integrated Genomics Operation. An average of 30-40 million paired reads was generated per sample. We used the log_2_foldchange > 1 or < -1 with *p_adj_*<0.05 as the threshold for significant changed genes. Quality control of FASTQ files was performed using the Rfastp R Bioconductor package (v0.1.2). Full genome sequence and transcript coordinates for the mm10 UCSC genomes and gene models were retrieved from the R Bioconductor packages BSgenome.Mmusculus.UCSC.mm10 (v1.4.0) and TxDb.Mmusculus.UCSC.mm10.knownGene (v3.4.0). Transcript abundance was determined from FASTQ files using Salmon (v0.8.1) [60], and transcript counts and TPM values were imported into R with the tximport R Bioconductor package (v1.8.0) [61]. Differential gene expression was performed with the DESeq2 R Bioconductor package (v1.20.0) [62]. For plots comparing the read counts of genes between samples, counts were normalized by dividing the raw read counts by the size factors for each sample as determined by DESeq2.

### rRNA-depletion RNA-seq and data analysis

Approximately 1 million cells were collected from 10 cm dishes. RNA extraction, rRNA depletion, library preparation, and RNA sequencing were performed by Novogene. An average of 30 million paired reads was generated per sample. Quantification of rRNA-depleted RNAseq reads over individual TE loci was performed using either the SQuIRE pipeline (v0.9.9.9a-beta - https://github.com/wyang17/SQuIRE) [46] or TE local (v1.1.1 - https://github.com/mhammell-laboratory/TElocal). The SQuIRE pipeline was used for alignment, counting, and calling of differential TEs. When using TElocal, reads were aligned with STAR (v2.27.10a) setting the ‘—winAnchorMultimapNmax’ and ‘—outFilterMultimapNmax’ arguments to 100[63], followed by counting with the TElocal function and differential abundance calculated with DESeq2 (v1.32.0). Reads were aligned to the mm10 genome sequence from the BSgenome.Mmusculus.UCSC.mm10 R Bioconductor package (v1.4.0).

### CUT&RUN

0.5 million cells from each line were collected. The detailed protocol for CUT&RUN for histone marks (H3K4me3, H3K27ac, H3K9me3, H3K27me3) and H3.3 are following previous described [34] with following modifications: final concentration of digitonin: 0.05%. Antibodies for CUT&RUN are following: H3.3 (Active Motif, 91191), H3K9me3 (Abcam, ab8898), H3K4me3 (Active Motif, 39159), H3K27me3 (Cell Signaling Technologies, #9733), H3K27ac (Active Motif, 39133), Rabbit IgG (Diagenode, C15410206). For ATRX CUT&RUN experiment, cells were fixed using 0.1% PFA in PBS for 1 minute at room temperature, followed by nuclei extraction with NE buffer (20 mM HEPES (pH7.9), 10 mM KCl, 0.1% Triton X-100, 20% Glycerol, 1 mM MnCl2, 1×cOmplete Mini-Tablet (1 tablet), 0.5 mM Spermidine) according to the protocol from EpiCypher (v2.0, https://www.epicypher.com/resources/protocols/cutana-cut-and-run-protocol/). The final concentration of digitonin buffer for ATRX CUT&RUN was 0.01%. For each sample, 2 µg ATRX antibody (Abcam, ab97508) was added. Libraries were prepared using the NEB Ultra II DNA library prep kit. Samples were PCR amplified for 14 cycles library pools was sequenced in the genomics core at the Rockefeller university.

### CUT&RUN alignment and differential peak analysis

Quality control of FASTQ files was performed using the Rfastp R Bioconductor package (v0.1.2). CUT&RUN reads were aligned using the Rsubread R Bioconductor package (v1.30.6), and predicted fragment lengths were calculated by the ChIPQC R Bioconductor package (v1.16.2) [64, 65]. The full mm10 genome sequence was retrieved from the R Bioconductor package BSgenome.Mmusculus.UCSC.mm10 (v1.4.0). Normalized, fragment-extended signal bigWigs were created using the rtracklayer R Bioconductor package (v1.40.6) [66]. Range-based heatmaps showing signal over genomic regions were generated using the profileplyr R Bioconductor package (v1.8.1) [67]. Any regions included in the ENCODE blacklisted regions of the genome were excluded from all region-specific analyses [68]. Bedgraph files generated with the deepTools package (v3.5.1) [69] were used for peak calling with SEACR in ‘stringent’ mode (v1.3) [70]. For H3K4me3, H3K27ac, H3K9me3, and H3.3, peaks were ranked based on the signal within peaks, and only the top 1% of peaks were used for downstream analysis. For ATRX, peaks were called compared to an IgG control sample.

Differential enrichment of signal within peaks was performed by counting the overlap of reads over a high confidence consensus peak set. This was obtained by reducing all replicates within all conditions to one peak set, and then keeping the peaks that overlap at least two out of the three replicates for any condition. Reads overlapping these peaks were counted for each replicate using the summarizeOverlaps function from the GenomicAlignments R Bioconductor package (v1.28.0) [71]. Differential peak enrichment between *Atrx* WT and *Atrx* KO replicates was then calculated using the DESeq2 R Bioconductor package (v1.32.0). To find differential enrichment in broad genomic regions for the H3K9me3 samples, reads were counted in 20 kb bins across the entire genome, and DESeq2 was used to find bins with enriched signal in the *Atrx* KO clones. Only bins in the top 10 percent in terms of read count were used for this analysis. The union of these differential bins and the differential SEACR peak regions was determined and used for downstream analysis of regions with enriched H3K9me3 signal in the *Atrx* KO clones.

Quantification of CUT&RUN signal over repeats at the individual locus and sub family level was performed with SQuIRE. The counts were transformed with rlog from the DESeq2 R Bioconductor package (v1.32.0), and these values were plotted on a heatmap using the ComplexHeatmap R Bioconductor package (v2.6.2) [72]. To assess ARTX CUT&RUN signal in telomeres, FASTQ files were aligned to a DNA sequence of 150 conserved telomere repeats (TTAGGG) using the R Bioconductor package Rbowtie2 (v1.12.0) [73, 74]. The z-scores of the log2(reads per million) were plotted on a heatmap using the ComplexHeatmap R Bioconductor package.

For quantification of ATRX CUT&RUN signal over genes, genes were divided into groups based on their expression levels. TPM values from the *Atrx* WT RNAseq dataset were calculated by Salmon and were split into five quantiles. The genomic location of each gene was then retrieved from the R Bioconductor package TxDb.Mmusculus.UCSC.mm10.knownGene (v3.10.0). IgG normalized bigwig signal for the ATRX CUT&RUN samples was obtained using the deepTools (v3.5.1) ‘bigwigCompare’ function, with the ‘operation’ argument set to ‘log2’. Signal over the gene regions (separated by RNAseq quantile) was computed with the deepTools computeMatrix function. The signal was visualized with the ggplot2 R package (v3.3.6).

### Omni-ATAC-seq

The samples are prepared according to the methods previous described [75-77] with minor modifications. 50,000 C3H/10T1/2 cells were collected for Omni-ATAC-seq. Amplification was performed with NEBNext High-Fidelity 2× PCR Master Mix (NEB, M0541s) and Nextera PCR Primers (8 cycles). The sequences of PCR primers with index adapters are listed in the Supplementary Table 31. The library pool was sequenced by Rockefeller University genomic core using a NextSeq High Output platform with 75bp paired-end reads in duplicates. An average of 30-40 million paired reads was generated per sample.

### ATAC-seq alignment and differential peak analysis

Quality control of FASTQ files was performed using the Rfastp R Bioconductor package (v0.1.2). ATAC-seq FASTQs were aligned to the mm10 genome from the Bsgenome.Mmusculus.UCSC.mm10 Bioconductor package (version 1.4.0) using Rsubread’s (v1.30.6) align method in paired-end mode with fragments between 1 to 5000 base-pairs considered properly paired [65]. Normalized, fragment signal bigWigs were created using the rtracklayer R Bioconductor package (v1.40.6) [66]. Peak calls for each replicate were made with MACS2 software in BAMPE mode [78].

Differential enrichment of signal within peaks was performed by counting the overlap of reads over a high confidence consensus peak set. This was obtained by reducing all replicates in all conditions to one peak set, and then keeping all peaks that overlap at least two out of the three replicates for any condition. Reads overlapping these peaks were counted for each replicate using the summarizeOverlaps function from the GenomicAlignments R Bioconductor package (v1.28.0) [71]. Differential peak enrichment between *Atrx* WT and *Atrx* KO replicates was then calculated using the DESeq2 R Bioconductor package (v1.28.1).

### Downstream peak analysis (CUT&RUN and ATAC-seq)

Peaks for both CUT&RUN and ATAC-seq were annotated with nearby genes using the rGREAT R Bioconductor package (v1.24.0) or with the closest gene and type of genomic region (e.g. promoter, intergenic, etc.) using the ChIPseeker R Bioconductor package (v1.28.3) [79-81]. The closest gene is defined as the gene for which the transcription start site is closest to the peak, and rGREAT defines a regulatory region around each gene and then annotates any peak in that region to that gene. For rGREAT annotation, the default settings were used to define regulatory regions of genes, which was 5Kb upstream and 1 kb downstream of each gene, and then extended a maximum of 1 Mb to the regulatory region of the next gene. Gene ontology analysis of genes associated with differential peaks (using the Biological Processes gene lists) was performed with the clusterProfiler R Bioconductor package (v4.0.5) [82]. Overlaps of differential peaks with other peak sets was done with the findOverlapsOfPeaks function from the ChIPpeakAnno R Bioconductor package (v3.26.4) [83]. Range heatmaps of CUT&RUN and ATAC-seq signal were made with the profileplyr R Bioconductor package [84]. For overlap of signal and peaks with enhancers, a peak file with enhancers from 10T cells was downloaded from the EnhancerAtlas 2.0 (http://www.enhanceratlas.org/). The downloaded peaks were converted from mm9 to mm10 using the ‘liftOver’ function from rtracklayer R Bioconductor package and the ‘mm9ToMm10.over.chain’ file downloaded from UCSC.

### ChromHMM analysis

Genome-wide ChromHMM (v1.23) models were generated using CUT&RUN signal for H3K4me3, H3K27ac, H3K27me3, H3K9me3, with IgG samples used as controls. Only the CUT&RUN samples from the *Atrx* WT cells were used to generate the models. Specifically, the BinarizeBam function was performed with default settings (mm10 genome), and this output was used in the LearnModel function. To determine the optimal number of states, we employed a previously published method to calculate the ratio of the ‘between sum of squares’ over the ‘total sum of squares’ of kmeans clustered emissions probabilities from all states of all models, considering 2-16 states [85]. The value of K at which the sum of squares ratio crossed 95% of the maximum was the 10 State model. To determine overlap of ChromHMM states with repeats, repeats from RepeatMasker were downloaded from UCSC (downloaded on Feb 4, 2022). To compare levels of overlap between states, the number of repeats that overlap a state was divided by the total number of regions in that state. A z-score of this value across all states was then plotted in a heatmap using the ComplexHeatmap R Bioconductor package (v2.6.2). A similar strategy was employed to measure ATRX peak overlap with ChromHMM states. H3K9me3 signal in specific states was quantified and visualized with a ranged heatmap using the profileplyr R Bioconductor package.

**Statistics**

Statistical analysis for proliferation, colony formation and qPCR were performed as indicated in the figure legends using paired or unpaired, two-tailed *t-test, or one-way ANOVA*. The analysis was not blinded.

## Data availability

All genomic and transcriptional data will be deposited in the Gene Expression Omnibus (GEO) at the time of publication.

## Supplementary Figures

**Figure S1:**
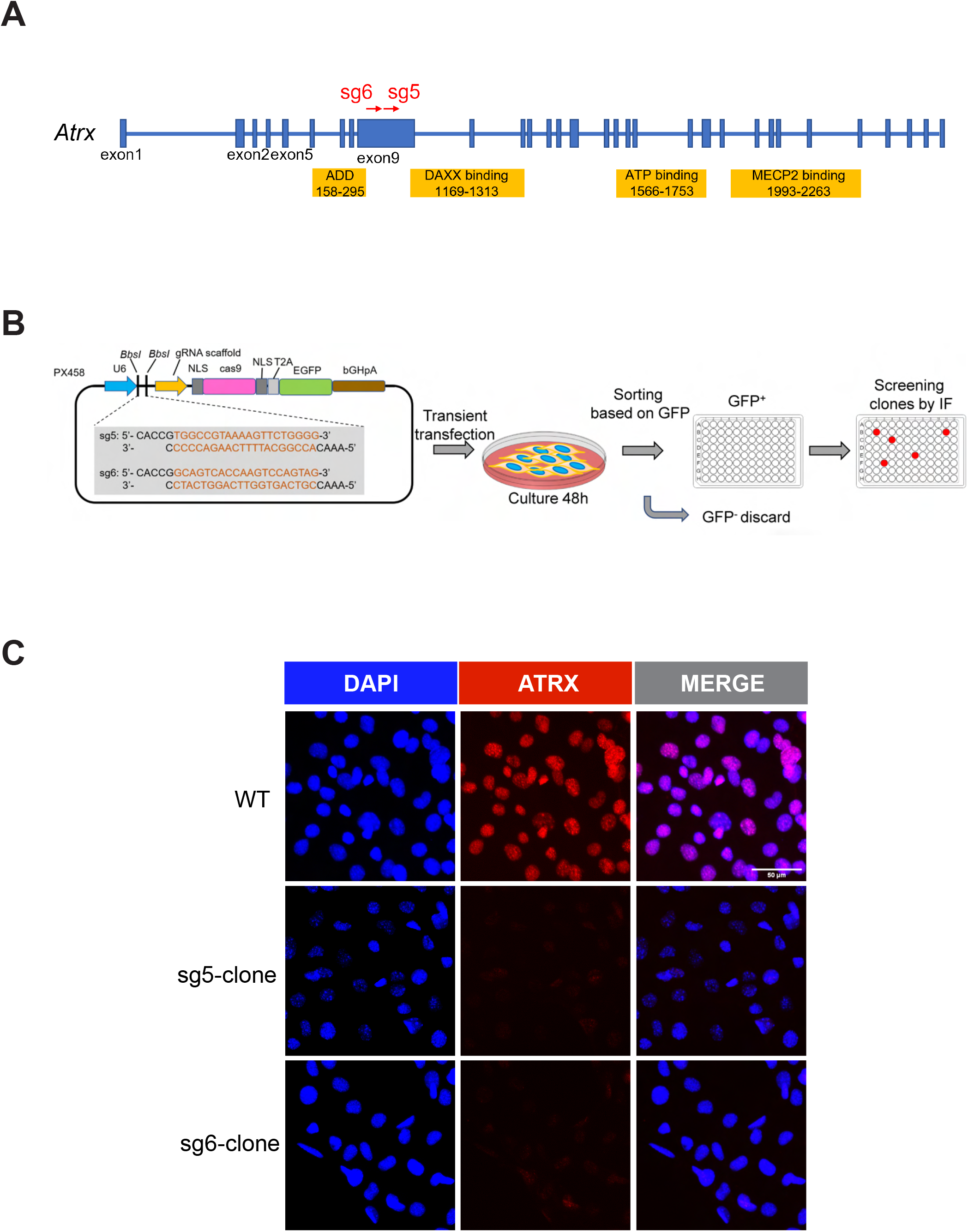
Generating ATRX knockout in C3H/10T1/2 cells. **(A)** Schematic of the of mouse *Atrx* gene structure. The yellow boxes indicate *Atrx* domains; blue boxes are the exons. The red arrows show the sgRNA target locations. **(B)** Workflow for establishing ATRX KO MPCs. The plasmid containing cas9 and sgRNAs targeting *Atrx* (WT control used an empty vector) was transiently transfected in cells. After 48 h, single cells expressing GFP were seeded in a 96 well plate (one cell per well). After 10 days, the cells were screened for ATRX expression (see panel **C**). Knockout clones from two different sgRNA were used in downstream experiments. **(C)** Immunofluorescence staining of ATRX in *Atrx* KO vs WT isogenic pairs. The scale bar is 50 µm.

**Figure S2:**
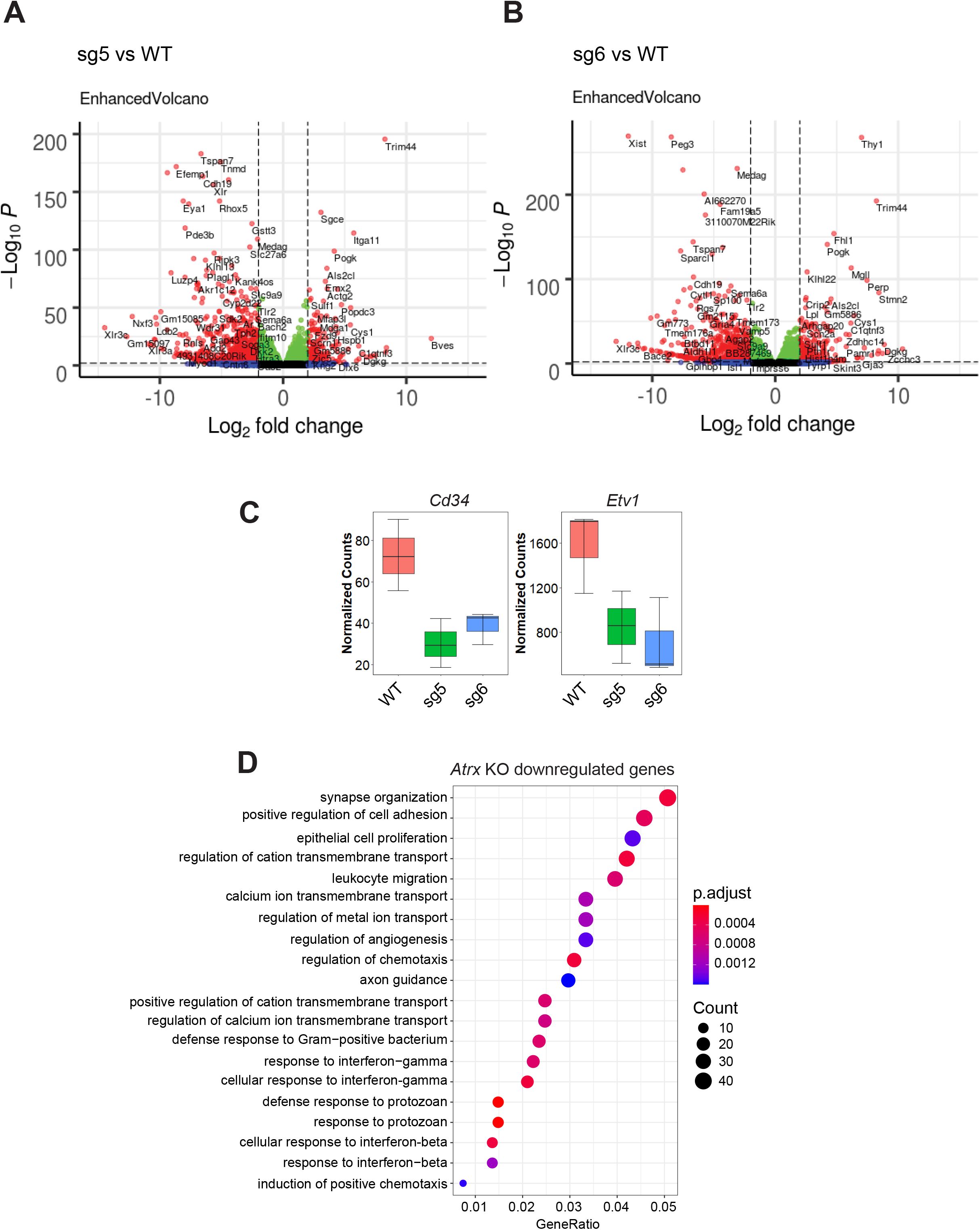
Loss of ATRX significantly perturbs the transcriptome of mesenchymal progenitor cells. **(A)** and **(B)** Volcano plots of each *Atrx* KO clone. The red points show genes which log_2_foldchange > 2 or < -2 and *p* <0.01. The blue points indicate genes with log_2_foldchange > 2 or < -2 and *p* value > 0.01. Green points indicate genes with log_2_foldchange between -2 to 2 and *p* <0.01. Black dots indicate genes with log_2_foldchange between -2 to 2 and *p* >0.01. **(C)** Boxplots for transcripts of interest. **(D)** GO analysis (biological process) for significantly down-regulated genes. Cutoffs used were log_2_foldchange < -1, and *p_adj_* < 0.05.

**Figure S3:**
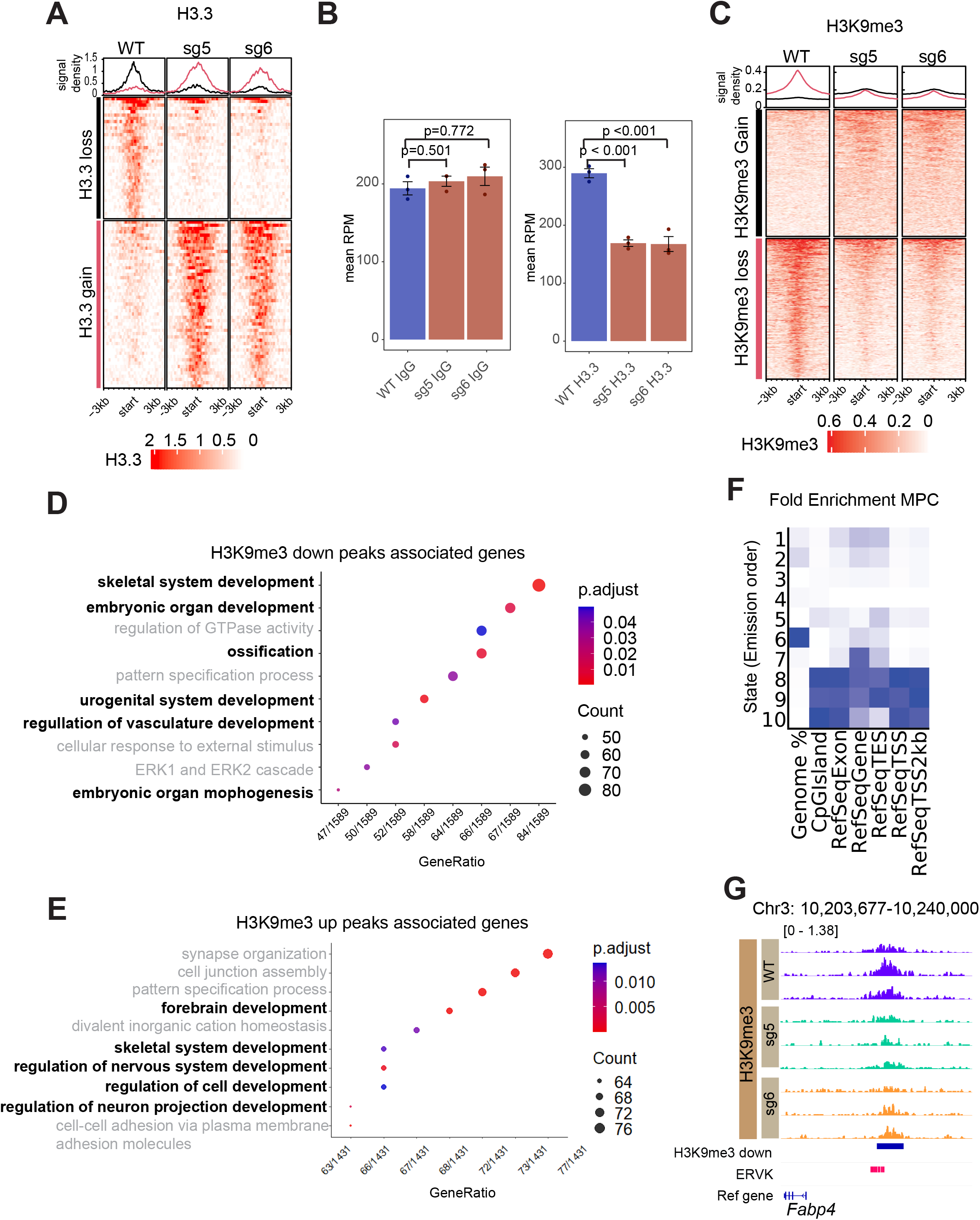
*Atrx* KO MPCs gain and loose H3.3 and H3K9me3 at specific loci. **(A)** Tornado plot of H3.3 signal at differential regions. **(B)** IgG and H3.3 signal at telomeric regions. RPM reads assigned per million mapped reads. The error bars show the standard deviations. The points show the data from individual biological replicates. *p* values were calculated using an unpaired one-way ANOVA with Tukey’s method for post hoc comparison. **(C)** H3K9me3 on differential regions. **(D)** The GO analysis (biological process) for significant (*p* ≤0.001) H3K9me3-lost regions or **(E)** H3K9me3-gained regions associated genes. **(F)** Average genome coverage and annotation of genic and non-genic elements in chromatin states determined by ChromHMM. **(G)** The Integrative genomic viewer tracks show H3K9me3 signals on *Fabp4* gene region. The blue bar shows the H3K9me3 significant down peaks (*p*-value < 0.01. The pink bar shows mouse ERVK elements.

**Figure S4:**
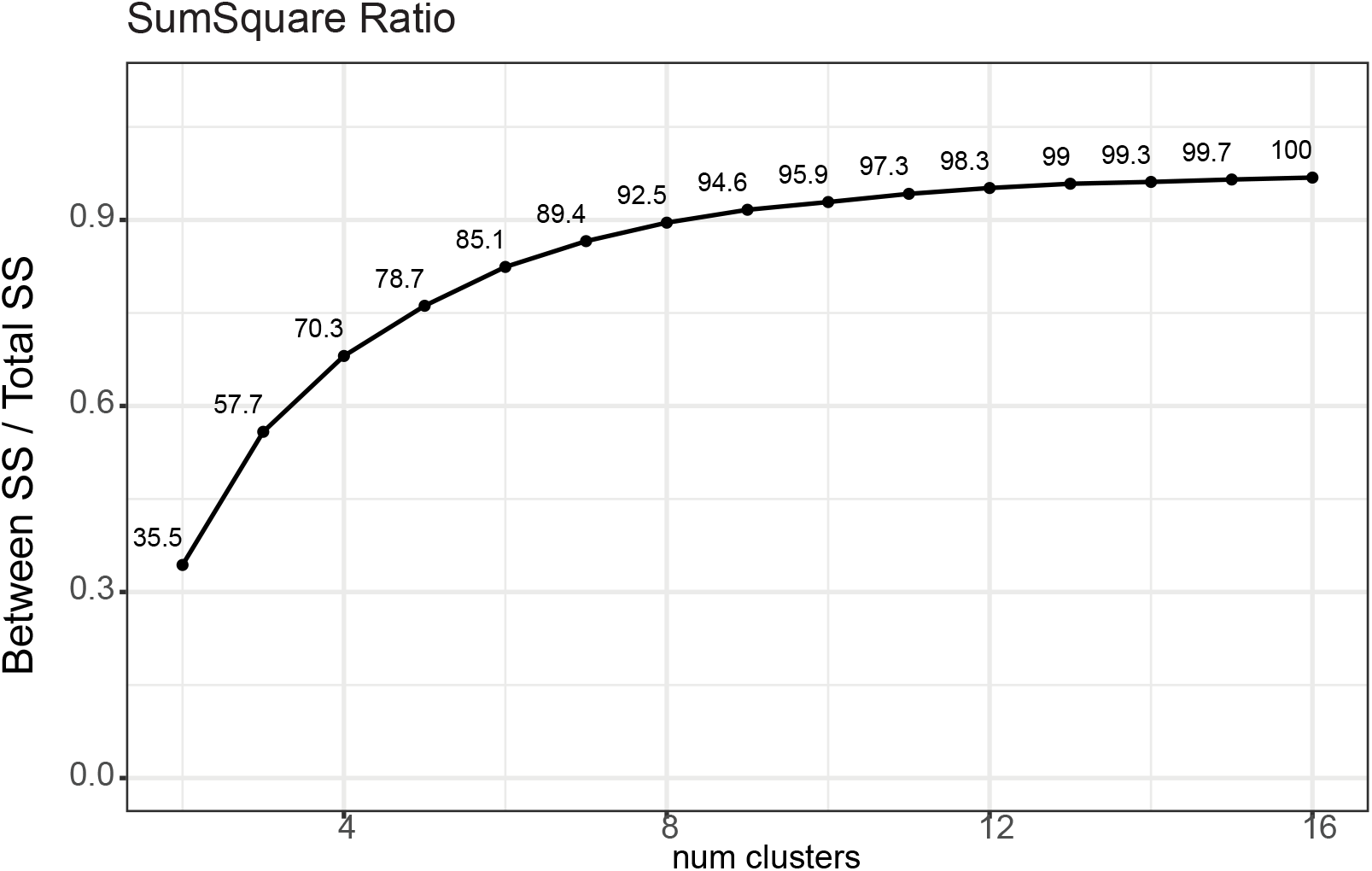
ChromHMM model. We established chromatin states using a multivariate hidden Markov model (ChromHMM) [36]. The plotted line demonstrates k-means clustering of the emission probabilities from *Atrx* WT model. We used the number of states that k-means equals to or higher than 95%. SS, sum of squares.

**Figure S5:**
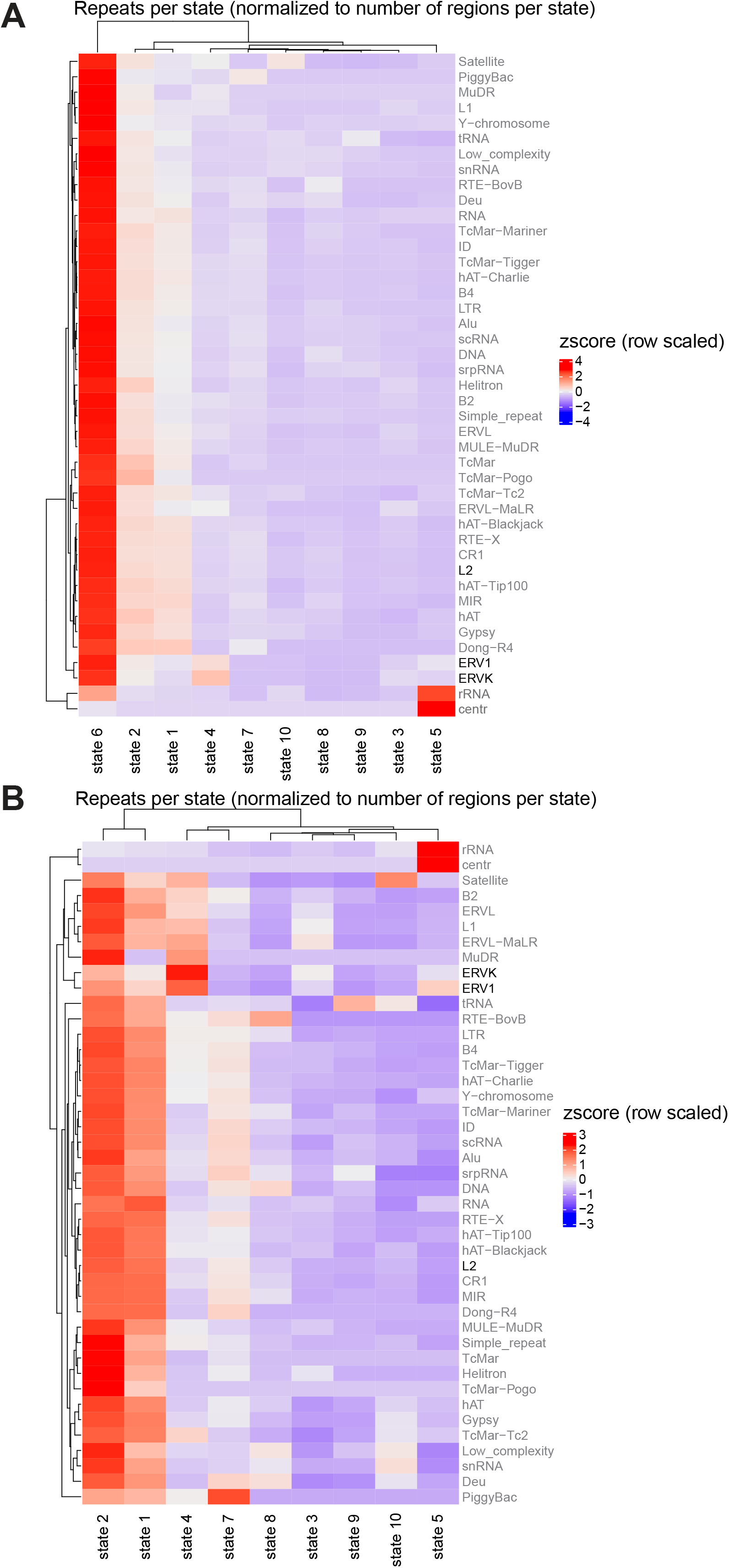
Enrichment of repetitive elements in chromatin states. **(A)** Heatmap of enrichment of repetitive elements in all ChromHMM-derived10 states. **(B)** Heatmap of enrichment of repetitive elements in 9 states excluding quiescent regions (state 6).

**Figure S6:**
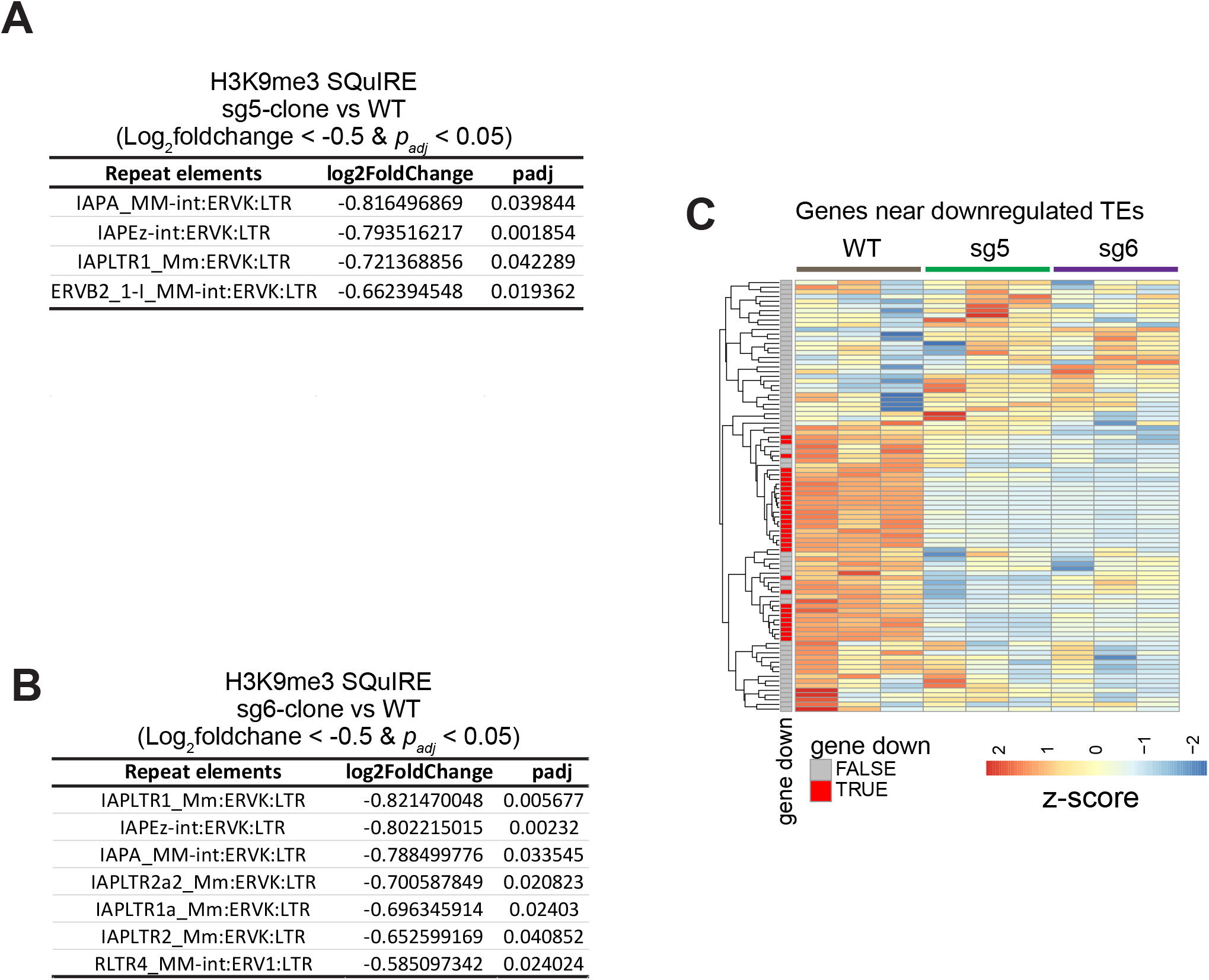
The association of H3K9me3 and TE expression. TEs associated with H3K9me3 loss (log_2_foldchange < -0.5 with *p_adj_* < 0.05) in each of two *Atrx* KO clones **(A)** and **(B)** vs WT MPCs. **(C)** The heatmap shows gene expression near downregulated TEs (TRUE: log_2_foldchange < -1 with *p_adj_* < 0.05).

**Figure S7:**
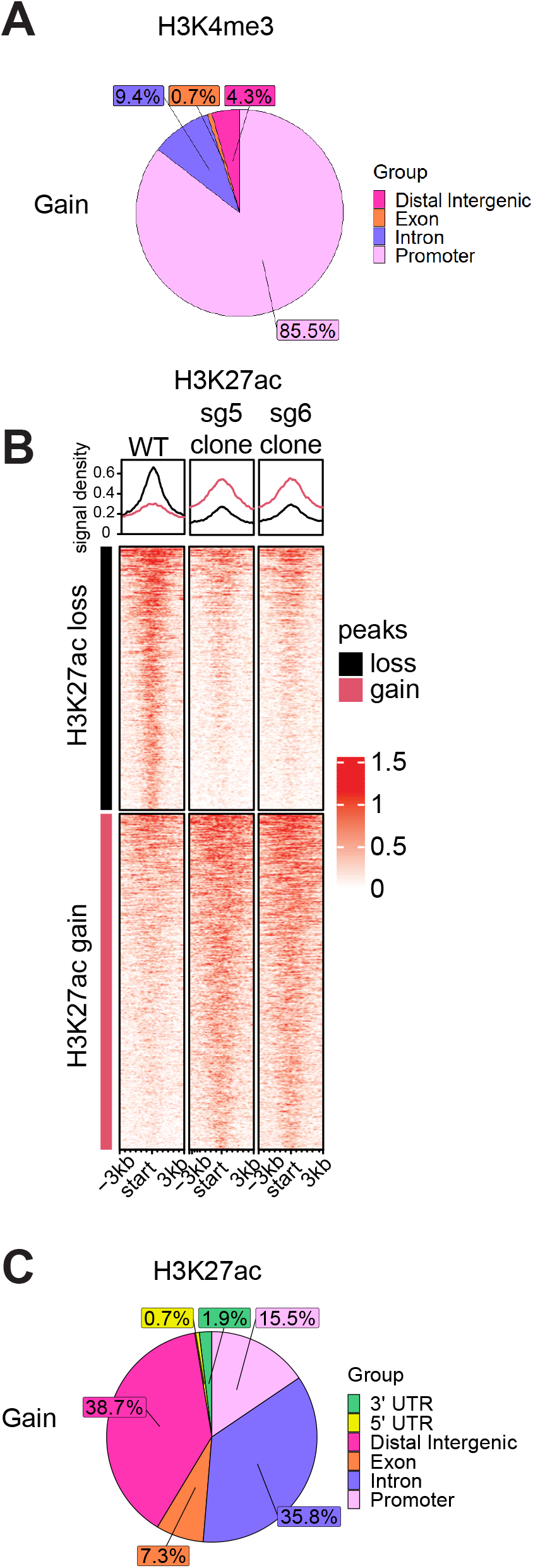
ATRX deficiency alters the distribution of active chromatin marks and chromatin accessibility. **(A)** Relative distribution of H3K4me3 gained peaks in *Atrx* KO MPCs by genomic feature. **(B)** Tornado plot of H3K27ac at differential regions. **(C)** Relative distribution of H3K27ac gained peaks in *Atrx* KO MPCs by genomic features.

**Figure S8:**
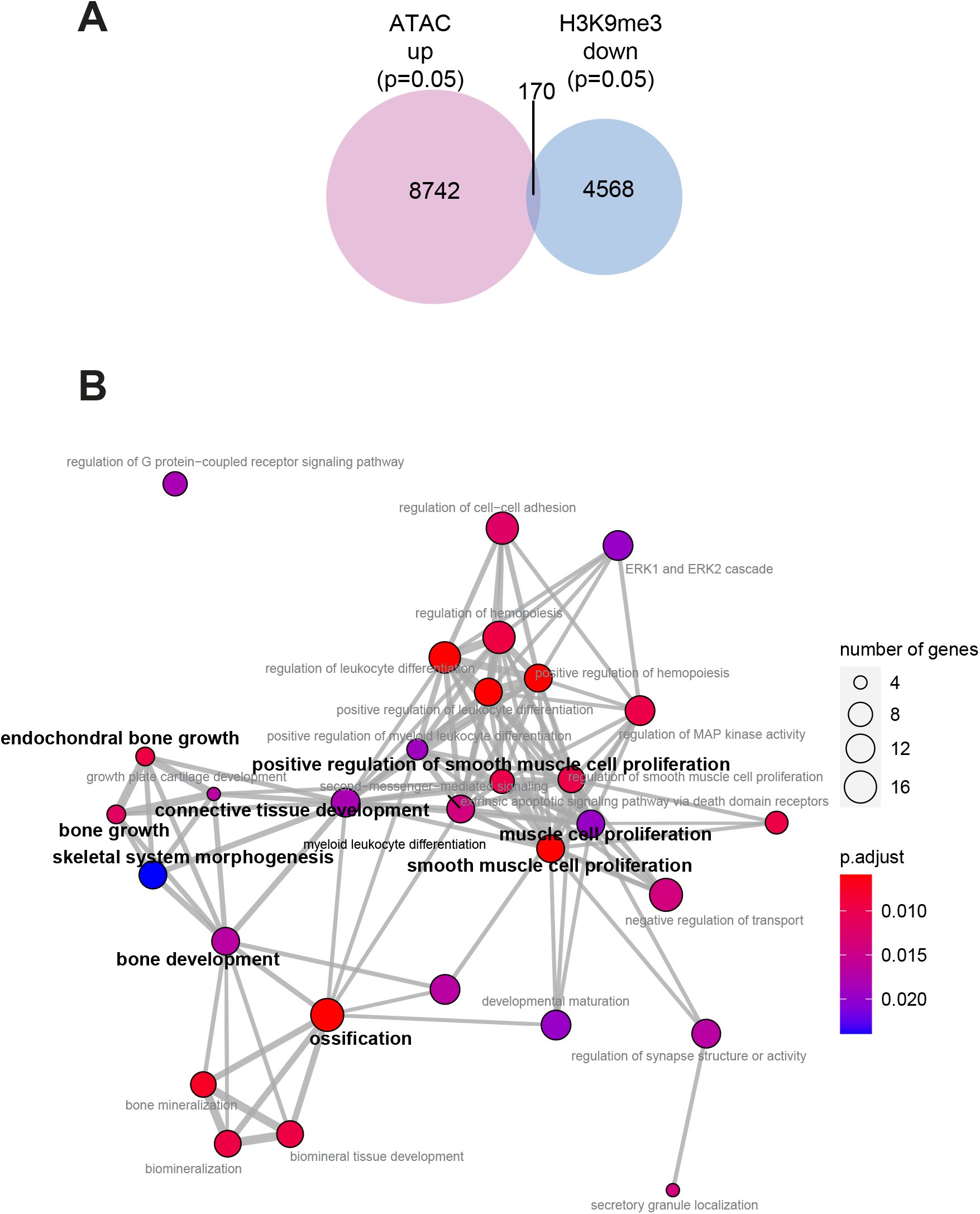
H3K9me3 depleted regions that gain chromatin accessibility are enriched in developmental genes in *Atrx* KO MPCs. **(A)** The intersection of ATAC-seq gained with lost H3K9me3 regions in *Atrx* KO versus WT MPCs. **(B)** Network plot of gene sets associated with the intersecting peaks in (A). Only significant (*p_adj_*<0.05) terms are shown.

**Figure S9:**
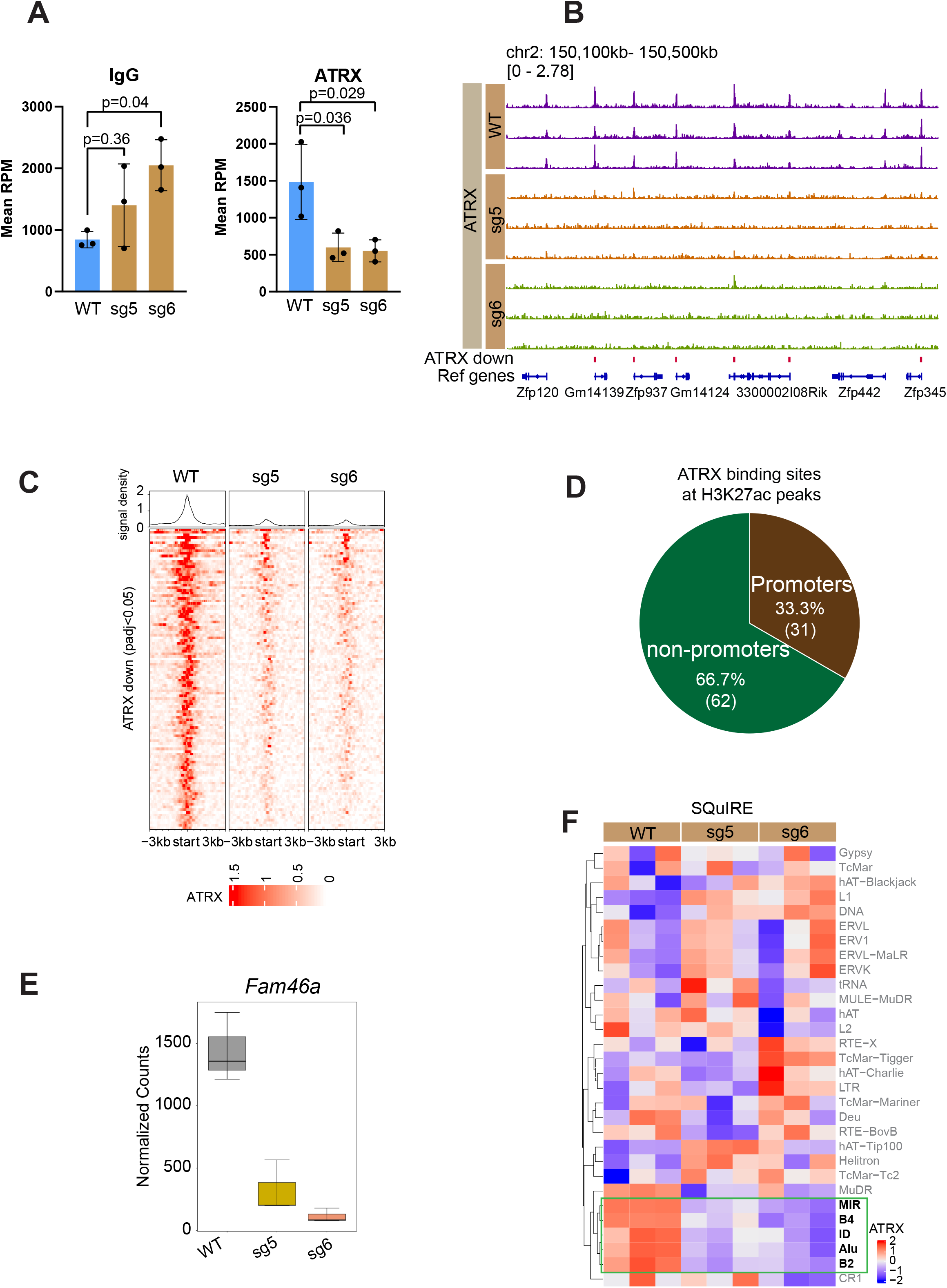
ATRX binds to repetitive elements in heterochromatin and to active regions in euchromatin. **(A)** ATRX and IgG signal at telomeric regions. The bar plots show IgG and ATRX signals on telomeres, respectively. The error bars show the standard deviations. The points show the data from individual biological replicates. *p* values were calculated using an unpaired one-way ANOVA with Tukey’s method for post hoc comparison. RPM (Reads per million mapped reads) assigned per million mapped reads. **(B)** Integrative genomic viewer tracks show typical regions of ATRX peaks (pink bars) at Zinc finger gene clusters. **(C)** ATRX signal at peak regions that have significantly higher signal in *Atrx* WT vs KO cells (*p_adj_* ≤0.05). **(D)** The percentage of ATRX binding sites at H3K27ac-enriched regions in *Atrx* WT cells. **(E)** Boxplot of normalized counts of *Fam46a* transcripts in MPC lines based on RNA-seq. **(F)** SQuIRE analysis shows that ATRX signals are reduced on specific repetitive elements.

---

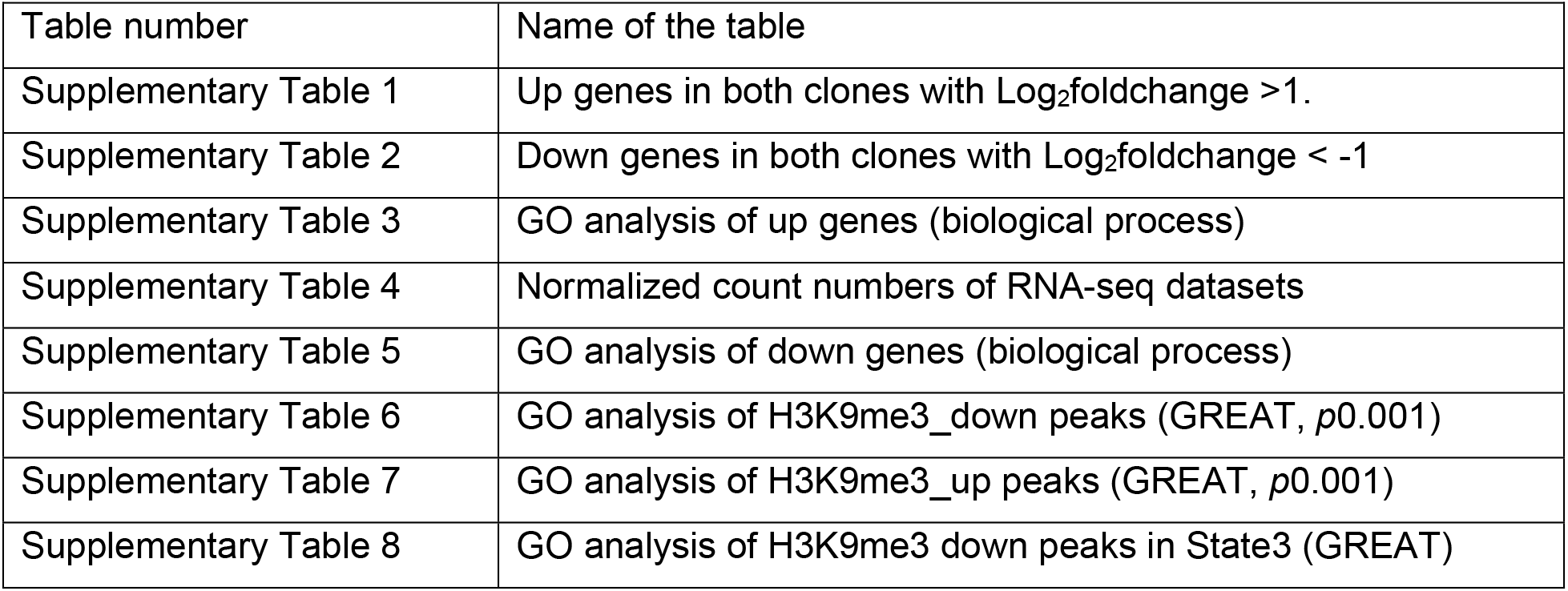
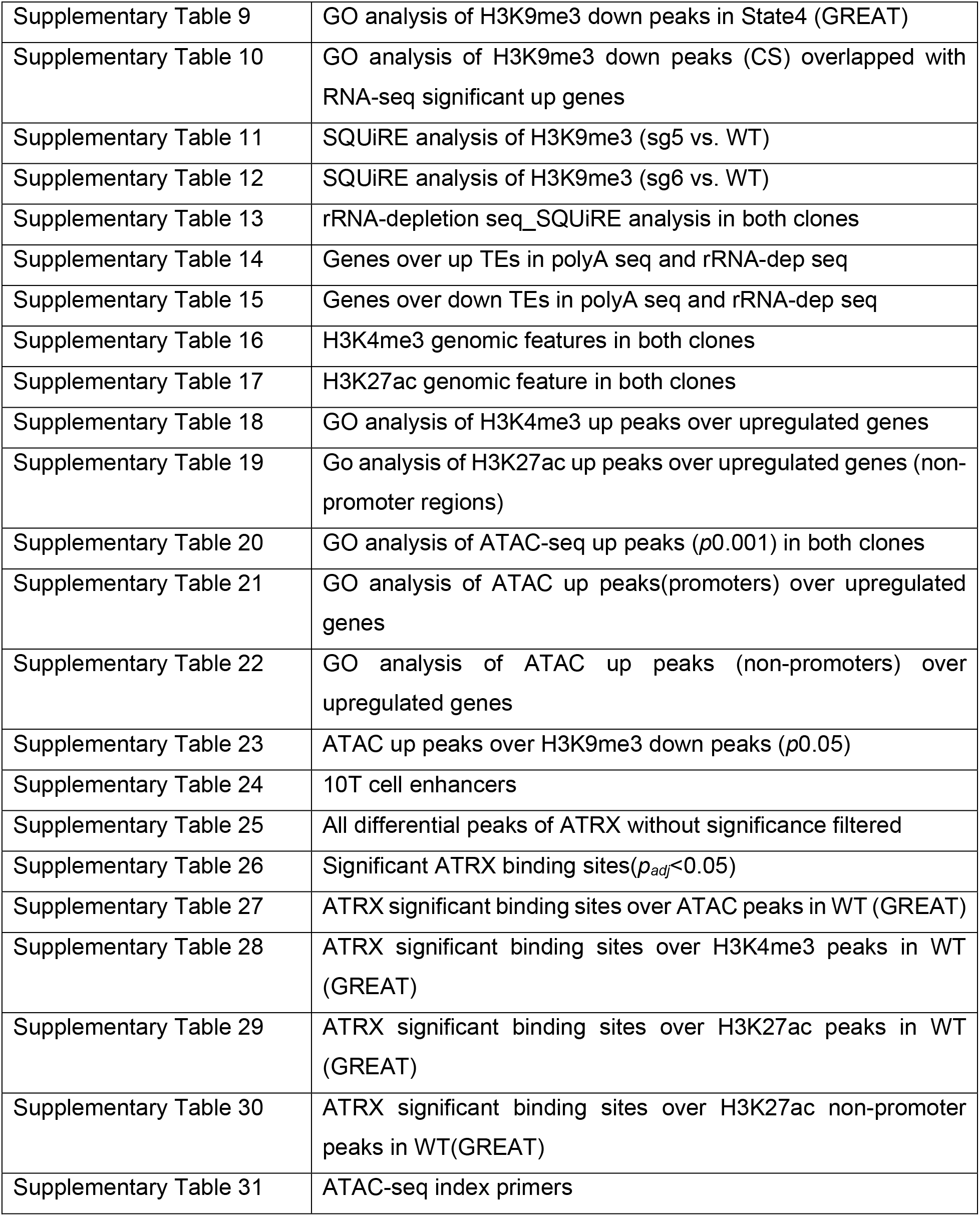
Tables.

